# Phylogenomics of the psychoactive mushroom genus *Psilocybe* and evolution of the psilocybin biosynthetic gene cluster

**DOI:** 10.1101/2022.12.13.520147

**Authors:** Alexander J Bradshaw, Virginia Ramírez-Cruz, Ali R. Awan, Giuliana Furci, Laura Guzmán-Dávalos, Paul Stamets, Bryn T.M. Dentinger

## Abstract

Psychoactive mushrooms in the genus *Psilocybe* have immense cultural value and have been used for centuries in Mesoamerica. Despite a recent surge in interest in these mushrooms due to emerging evidence that psilocybin, the main psychoactive compound, is a promising therapeutic for a variety of mental illnesses, their phylogeny and taxonomy remain substantially incomplete. Moreover, the recent elucidation of the psilocybin biosynthetic gene cluster is known for only five species of *Psilocybe*, four of which belong to only one of two major clades. We set out to improve the phylogeny for *Psilocybe* using shotgun sequencing of 71 fungarium specimens, including 23 types, and conducting phylogenomic analysis using 2,983 single-copy gene families to generate a fully supported phylogeny. Molecular clock analysis suggests the stem lineage arose ∼66 mya and diversified ∼53 mya. We also show that psilocybin biosynthesis first arose in *Psilocybe*, with 4–5 possible horizontal transfers to other mushrooms between 40 and 22 mya. Moreover, predicted orthologs of the psilocybin biosynthetic genes revealed two distinct gene orders within the cluster that corresponds to a deep split within the genus, possibly consistent with the independent acquisition of the cluster. This novel insight may predict differences in chemistry between the two major clades of the genus, providing further resources for the development of novel therapeutics.

## Introduction

Members of the renowned psychoactive genus *Psilocybe* (Fr.) P. Kumm., colloquially known as “magic mushrooms,” have become a major scientific and public curiosity since Robert Gordon Wasson first published his infamous *Life* magazine piece “Seeking the Magic Mushroom” in 1957 (Wasson 1957)*. Psilocybe* sensu lato is a globally distributed genus of psychoactive mushrooms and in its traditional concept contains between 277 and 326 known species (Guzmán 2005; Kirk et al. 2008). Molecular phylogenetic studies have demonstrated that *Psilocybe* s.l. is not monophyletic (Moncalvo et al. 2002; Matheny et al. 2006). One group consists of species that exhibit a “bluing” reaction when damaged, a feature that is thought to indicate the presence of the psychoactive alkaloid psilocybin (Guzmán 2005; Lenz et al. 2020, 2021). However, only 24 of ∼144 species with the bluing reaction have been included in multi-locus molecular phylogenetic datasets (Borovička et al. 2011; Ramírez-Cruz et al. 2013a).

While taxonomy of *Psilocybe* has been the subject of a handful of studies spanning the past 60+ years (Singer and Smith 1958; Hofmann et al. 1959; Guzmán 1983, 1995, 2005; Borovička et al. 2011; Ramírez-Cruz et al. 2013b; Cortés-Pérez et al. 2020), the ambiguous legal status of *Psilocybe* specimens and their rampant misidentification has hindered modern taxonomic research on them (Bradshaw et al. 2022). The only multi-locus molecular phylogenetic studies included 24 (accounting for found synonyms) species and up to three gene regions, with many branches without strong statistical support and lacking most of the currently recognized species (Borovička et al. 2011, 2015; Ramírez-Cruz 2013a). Moreover, most of the available molecular data do not come from type specimens, making it impossible to authenticate modern material. Together, our poor phylogenetic understanding and uncertainty in the application of names limit our ability to investigate evolutionary questions in *Psilocybe* and impair the construction of a robust predictive framework for exploring genetic traits of potential interest for the development of novel therapeutics.

The enzymatic synthesis of the psychoactive compound psilocybin was first elucidated in *Psilocybe cubensis* (Earle) Singer and *P. serbica* M.M. Moser & E. Horak (Fricke et al. 2017). Four core genes encoding enzymes that convert the amino acid tryptophan into psilocybin occur in a cluster: PsiD (tryptophan decarboxylase), PsiK (kinase), PsiM (methyltransferase), and PsiH (P_450_) (Fricke et al. 2017). Phylogenetic analysis indicated that this cluster has been horizontally acquired by psilocybin-producing mushrooms in other genera with common ecological niches (Awan et al. 2018; Reynolds et al. 2018), although the direction and relative timing of these transfers remains unclear. Notably, only a select few species of *Psilocybe* have been investigated at a genomic scale (Fricke et al. 2017; Awan et al. 2018; Reynolds et al. 2018; McKernan et al. 2021; Dörner et al. 2022), which does not represent all of the known ecologies, and they do not span the phylogenetic range of the two major clades identified by Ramírez-Cruz et al. (2013a), severely limiting inferences of evolutionary patterns.

Due to their relatively small genomes, whole genome sequencing of Fungi has been advancing at an expedited pace (Grigoriev et al. 2014). One of the major challenges to fungal molecular systematics has been the lack of available material due to the unpredictable and ephemeral nature of fungal reproductive structures, and the poor preservation of DNA in fungarium specimens. However, the technical impediment to utilizing preserved specimens has been largely overcome and it is now feasible and relatively cost-effective to generate whole metagenomes from fungarium specimens for phylogenomics (Dentinger et al. 2016 Tremble et al. 2020, Liimatainen et al. 2022). In this study, we aimed to vastly improve our understanding of *Psilocybe* evolution by increasing the species represented in a genome-scale phylogenetic dataset by exploiting fungarium resources and using the phylogeny as a backbone constraint to incorporate all publicly available DNA sequence data for *Psilocybe* and to explore the evolution of psilocybin biosynthesis.

## Results

### Sequencing and genome assembly

Genomic DNA (gDNA) libraries were prepared using Nextera DNA Flex Library Prep and sequenced using Novaseq 2x 150bp. Total reads for samples ranged from 3,909,488 reads (Psilocybe_mexicana_IBUG-13593, ∼11.7x coverage) to 225,748,454 reads (Psilocybe_baeocystis_WTU-F-011245, ∼ 677.2x coverage). Across all of our samples, we achieved an average of 94,490,694 reads (∼283.5x coverage) and a median of 99,070,482 reads (∼297.2x coverage).

Following genome assembly, statistics for each sample were measured with N50 values ranging from 554 (Psilocybe_tuberosa_WTU-F-011378) to 60,042 (Psilocybe_stuntzii_WTU-F- 011520) and the number of contigs for each assembly ranging from 7,004 (Psilocybe_stuntzii_WTU-F-011520) to 784,732 (Psilocybe_caerulescens_var_mazatecorum_SFSU-F-029971). It should be noted that these assemblies are likely to be highly non-contiguous due to the fact they are from museum vouchers, which are susceptible to high amounts of contamination and should be treated as metagenome assemblies (Dentinger et al. 2016).

In addition to looking at assembly statistics, we compared the recovery of single-copy orthologs based on the agaricales_odb10 reference database to judge the “completeness” of our genomes. We observed BUSCO scores ranging from 30.7% (Psilocybe_fuliginosa_NY-1901148) to 95.4% (Psilocybe_stuntzii_WTU-F-011520). In terms of complete BUSCO gene recovery, only one sample achieved a score of ∼95% (the general measurement of a high-quality assembly), yet phylogenomic data utilizing museum vouchers have been shown to allow for robust generation of phylogenetic relationships with strong support values, especially those at deeper nodes, which are often difficult to resolve (Dentinger et al. 2016) All assembly statistics and BUSCO scores for each sample are reported in Supplementary Table 1.

### *Psilocybe* phylogenomic species tree and divergence time of major clades

Specimens of *Psilocybe* were selected based on species-level voucher designation and totaled 71 specimens, 23 of which were type specimens, the ultimate name-bearing specimen for a species (Aime et al. 2021) (Table 1). Of the 71 specimens, 52 were true *Psilocybe* (*Psilocybe* sensu stricto) (20 type specimens) (Figure 1), 19 were found to not be true *Psilocybe*: 14 specimens clustered together phylogenetically, and belonged to *Deconica* (W.G. Sm.) P. Karst. (*Psilocybe* sensu lato) (3 type specimens), 3 clustered together, only identifiable to *Strophariaceae,* one specimen was found to b*e Kuehneromyces* sp. (Psilocybe_laticystis_UBC-F16759), and one sample was unable to be accurately identified (Psilocybe_washingtonensis_WTU-F-055019) and was removed from the further downstream analysis (Supplementary Figure 1, Table 1).

**Figure 1:**
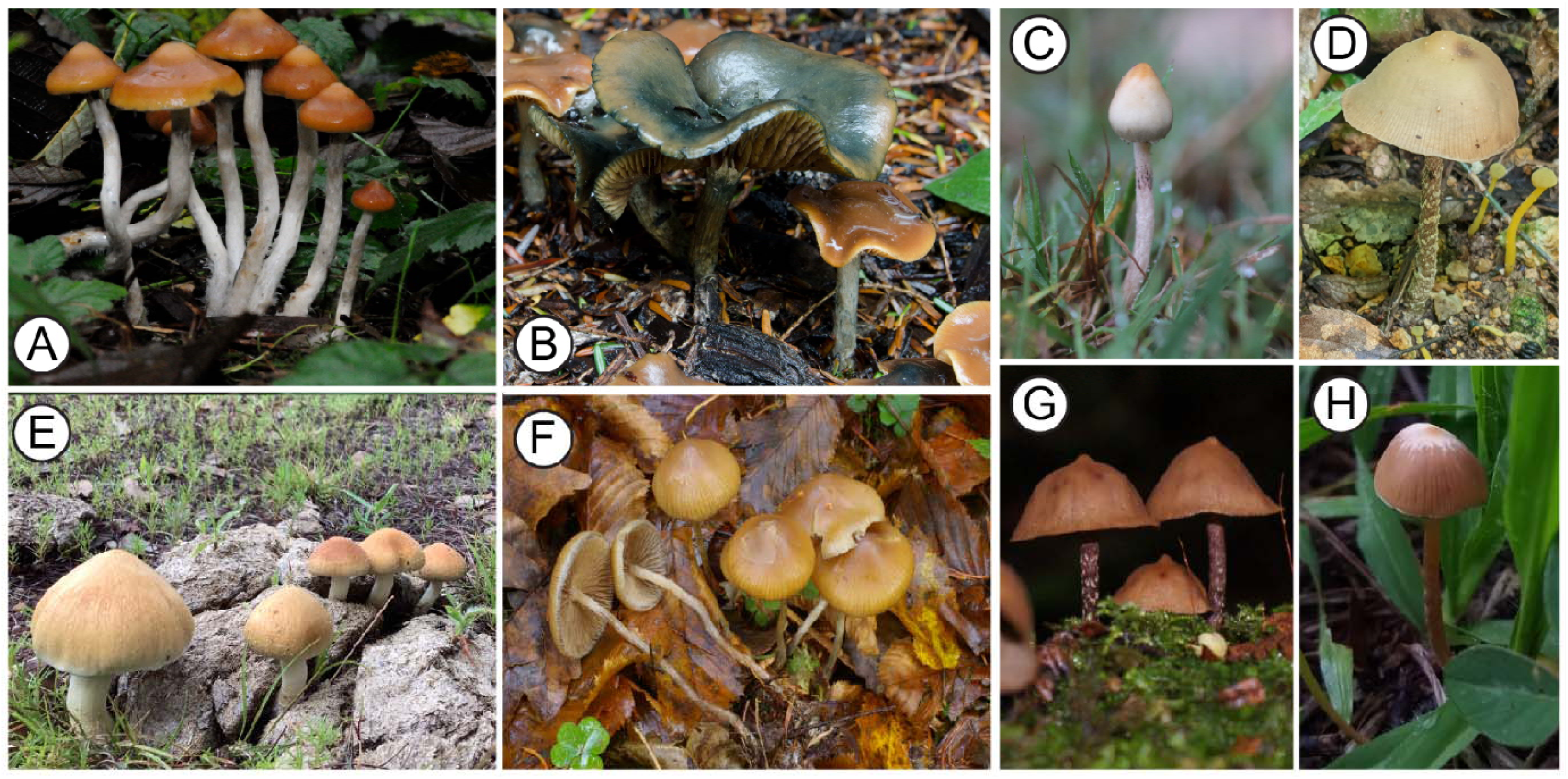
Pictorial representation of *Psilocybe* species.

**A**, *Psilocybe azurescens* (image credit, Paul Stamets), **B**, *Psilocybe cyanescens* (image credit Bryn T.M. Dentinger), **C**, *Psilocybe semilanceat*a (image credit Paul Stamets), **D**, *Psilocybe zapotecorum* (image credit Bryn T.M. Dentinger), **E,** *Psilocybe cubensis* (image credit Oscar Castro-Jauregui), **F**, *Psilocybe bohemica* (image credit Jan Borovička), **G**, *Psilocybe yungensis* (image credit Virginia Ramírez-Cruz), **H**, *Psilocybe mexicana* (image credit Oscar Castro- Jauregui).

**Table 1:**
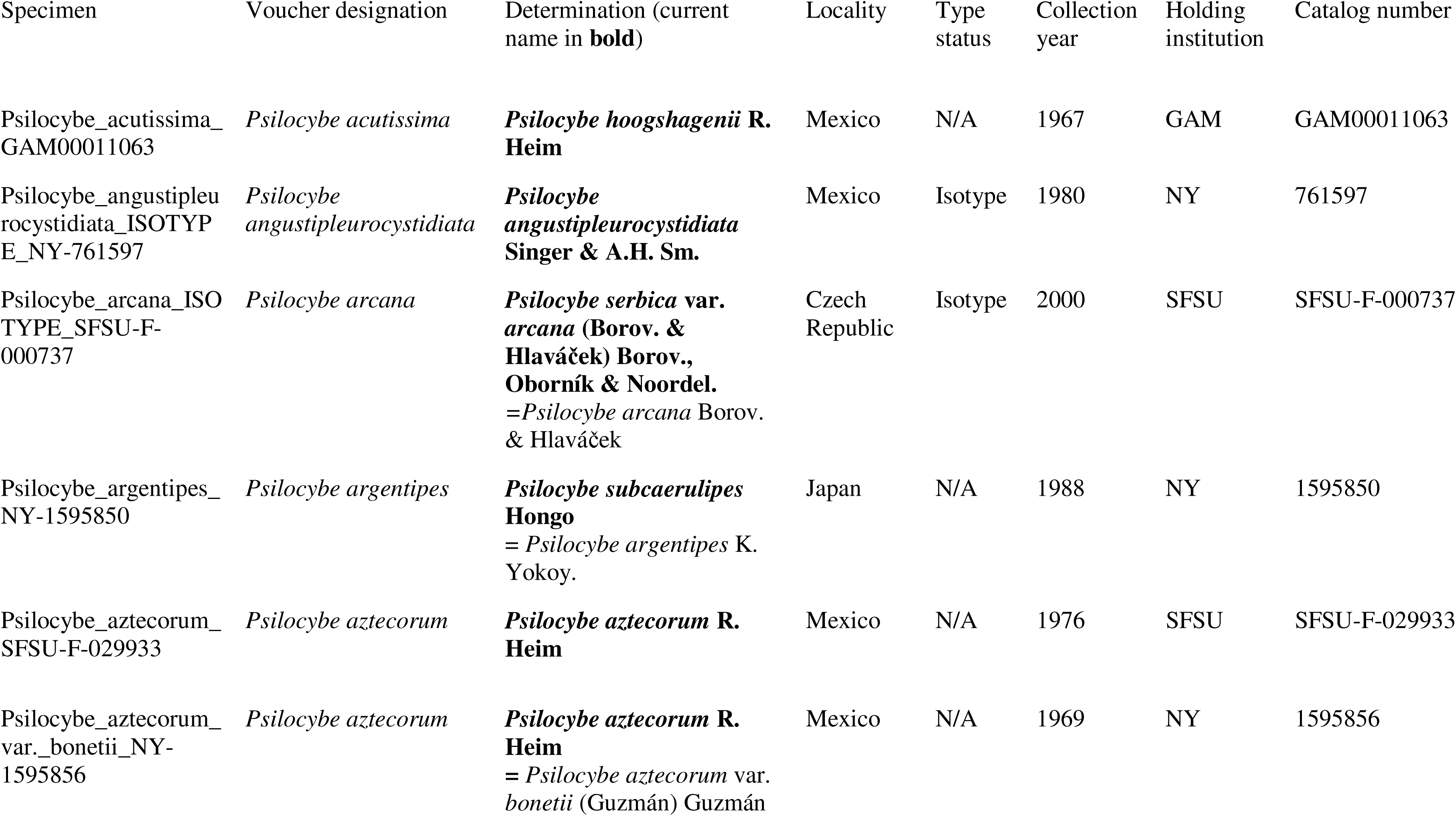
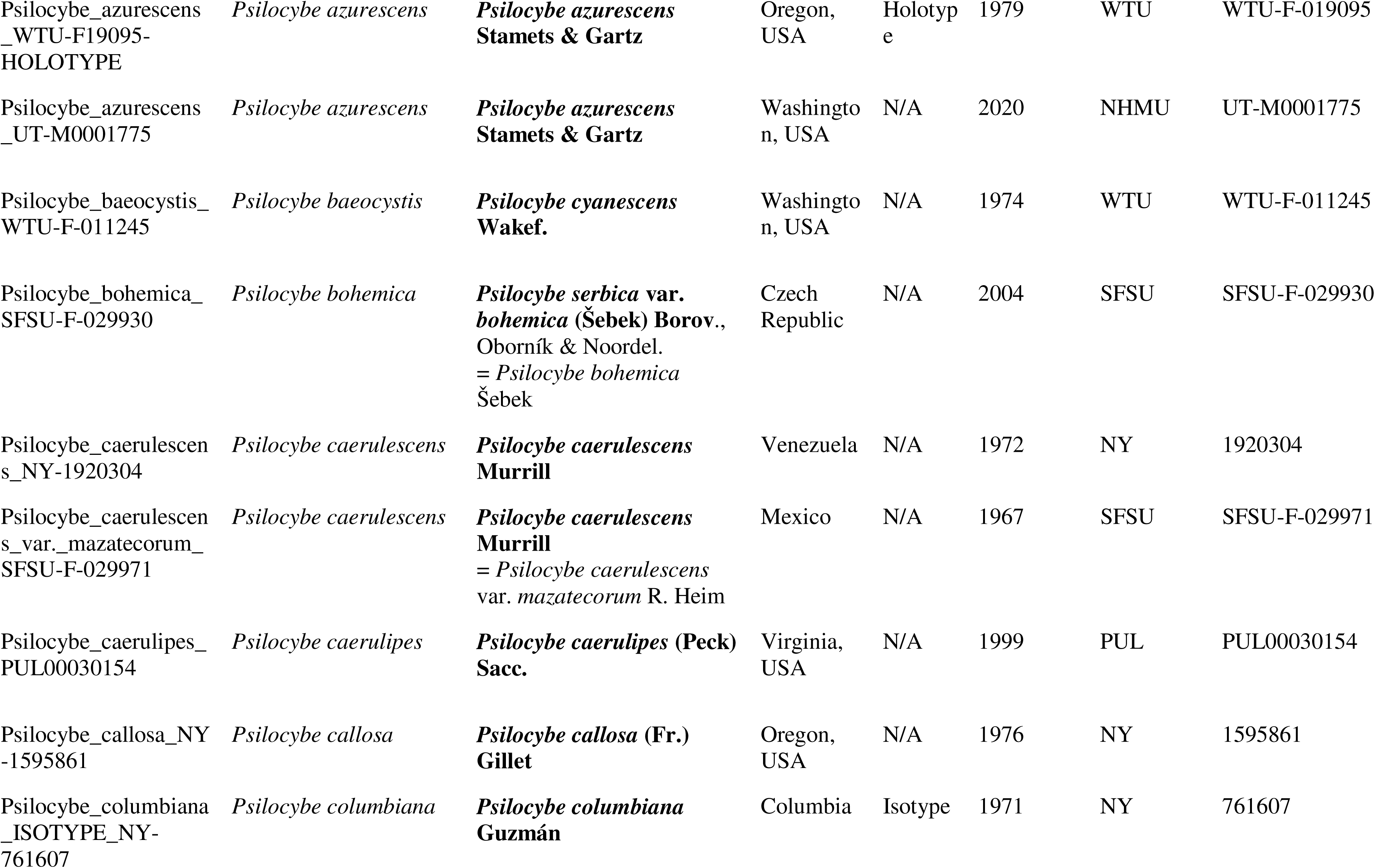
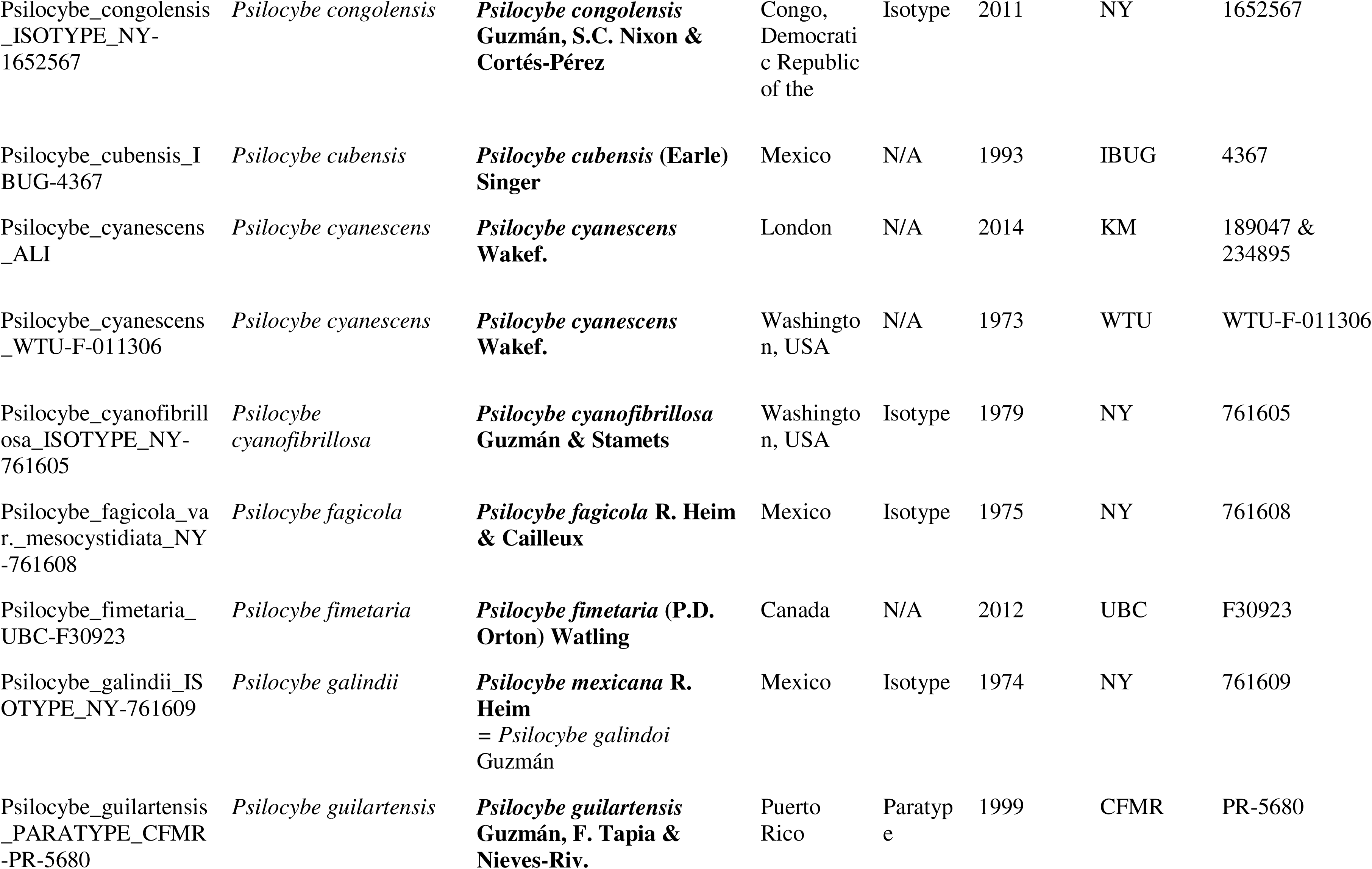
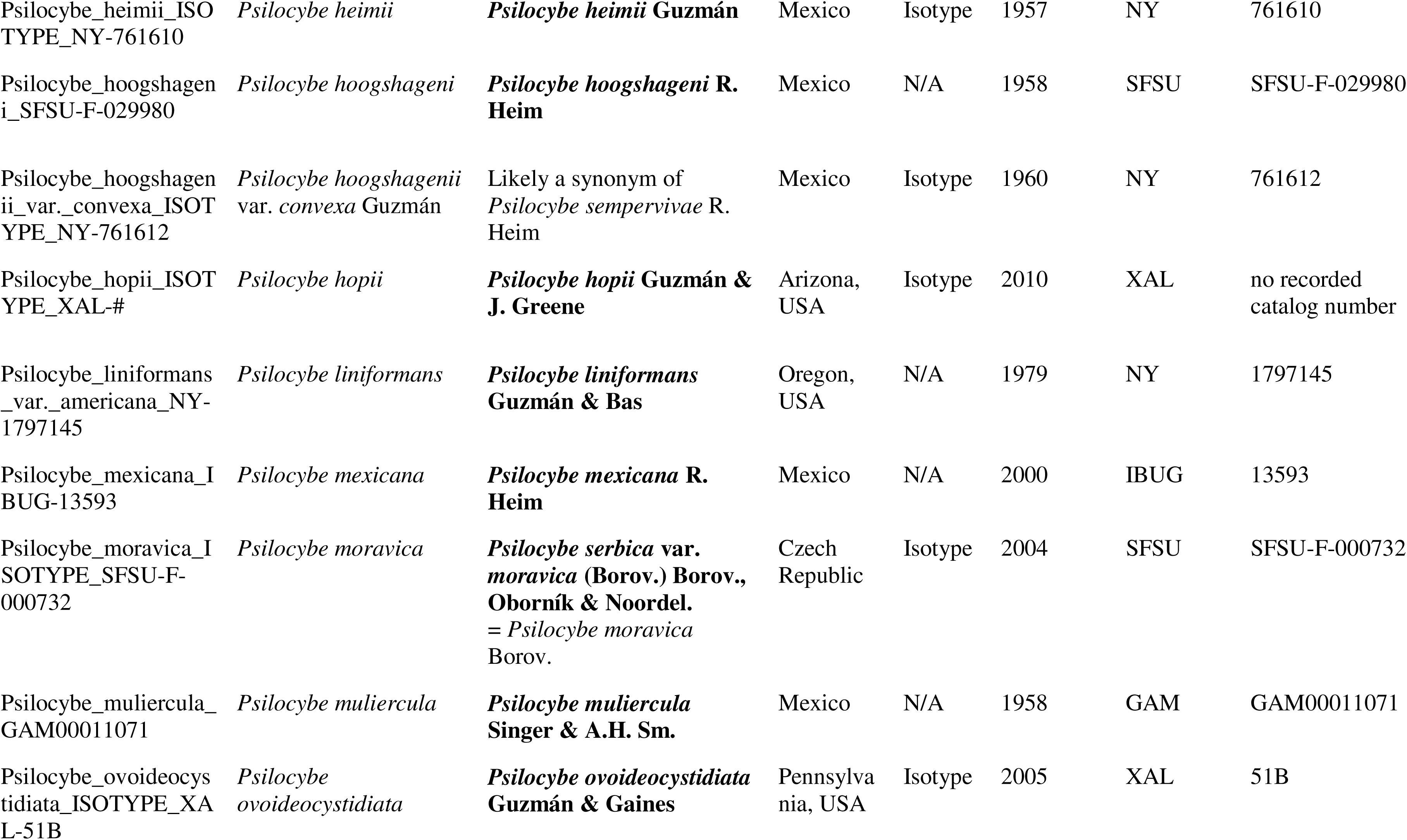
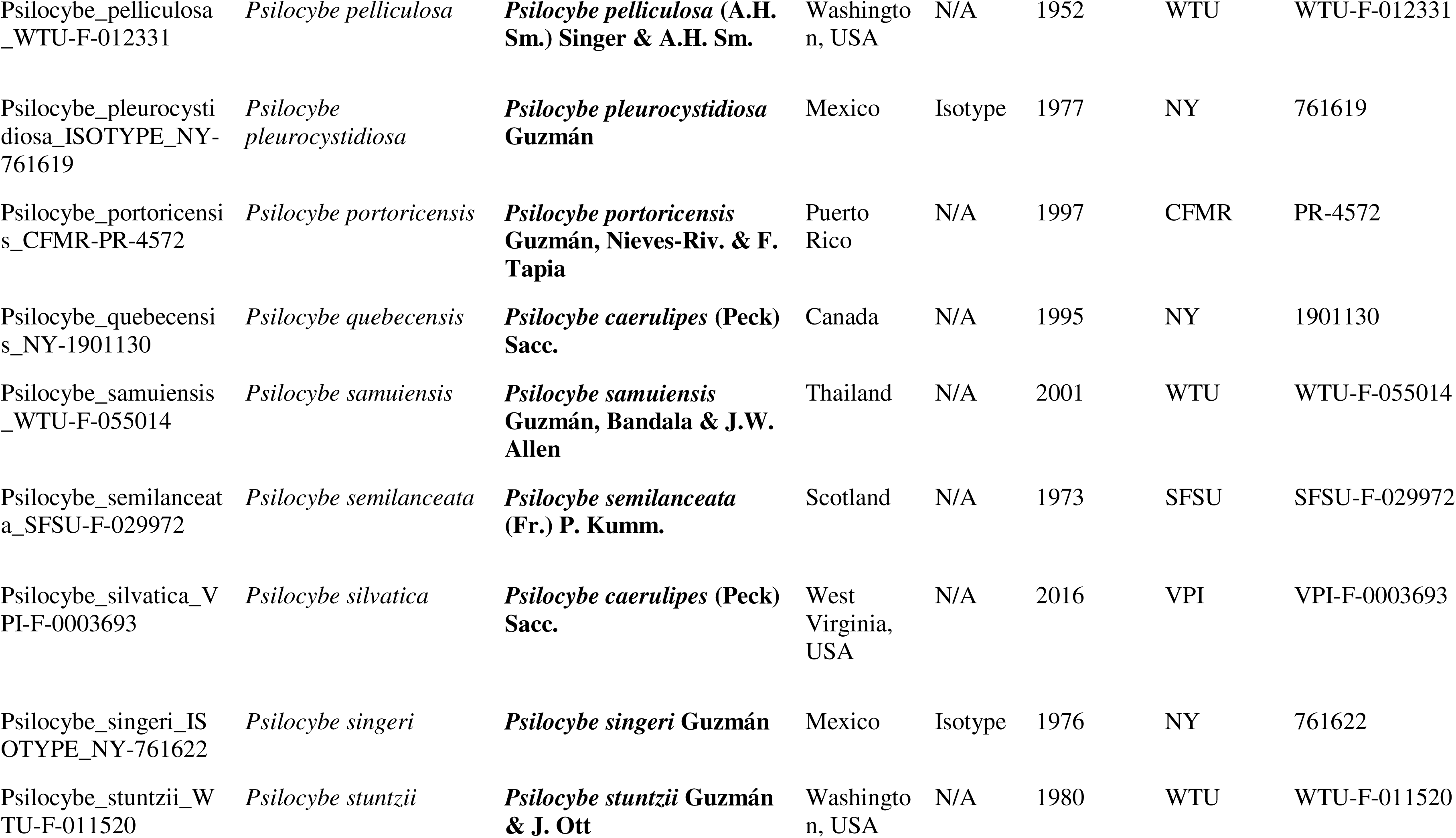
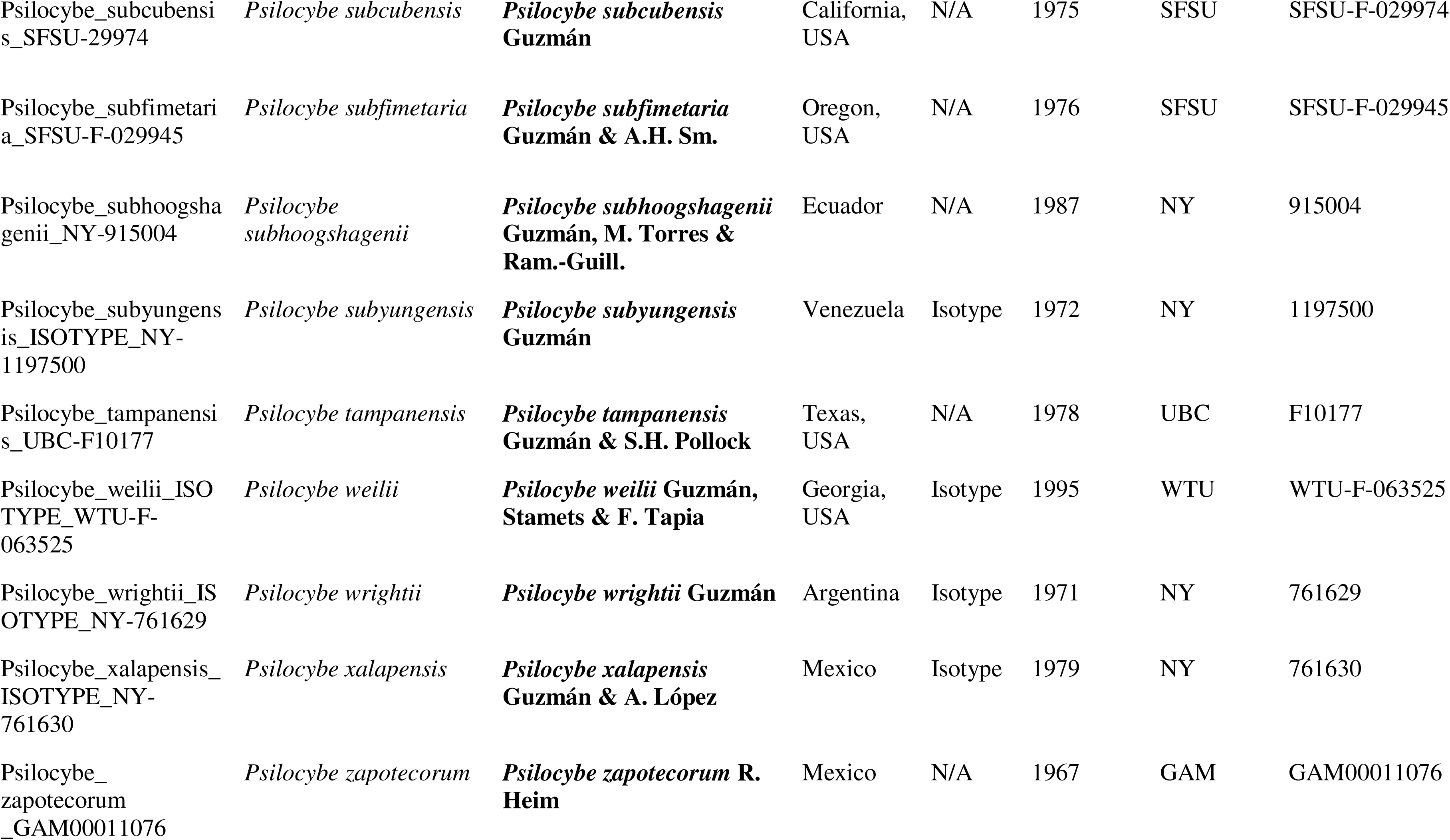
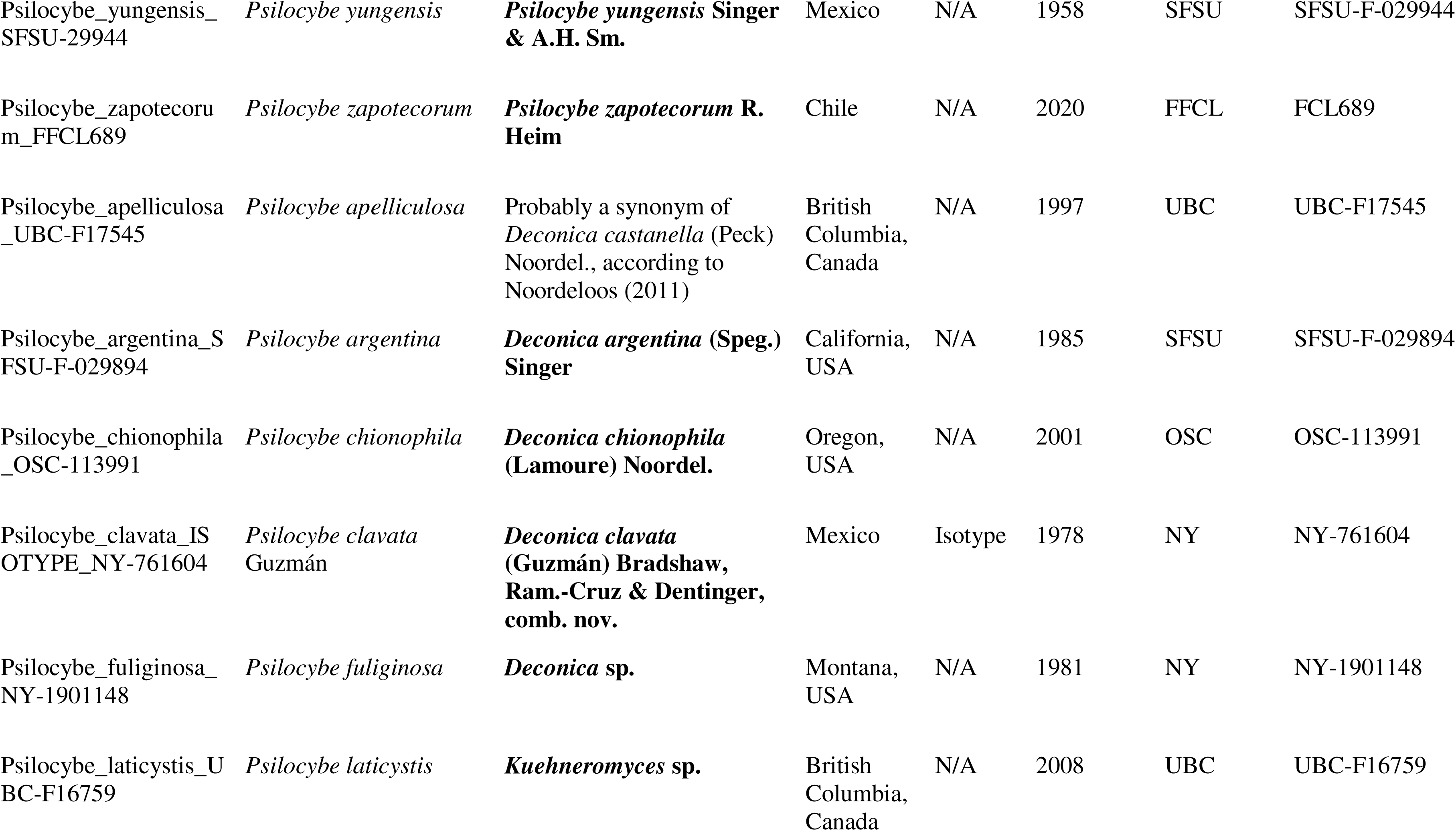
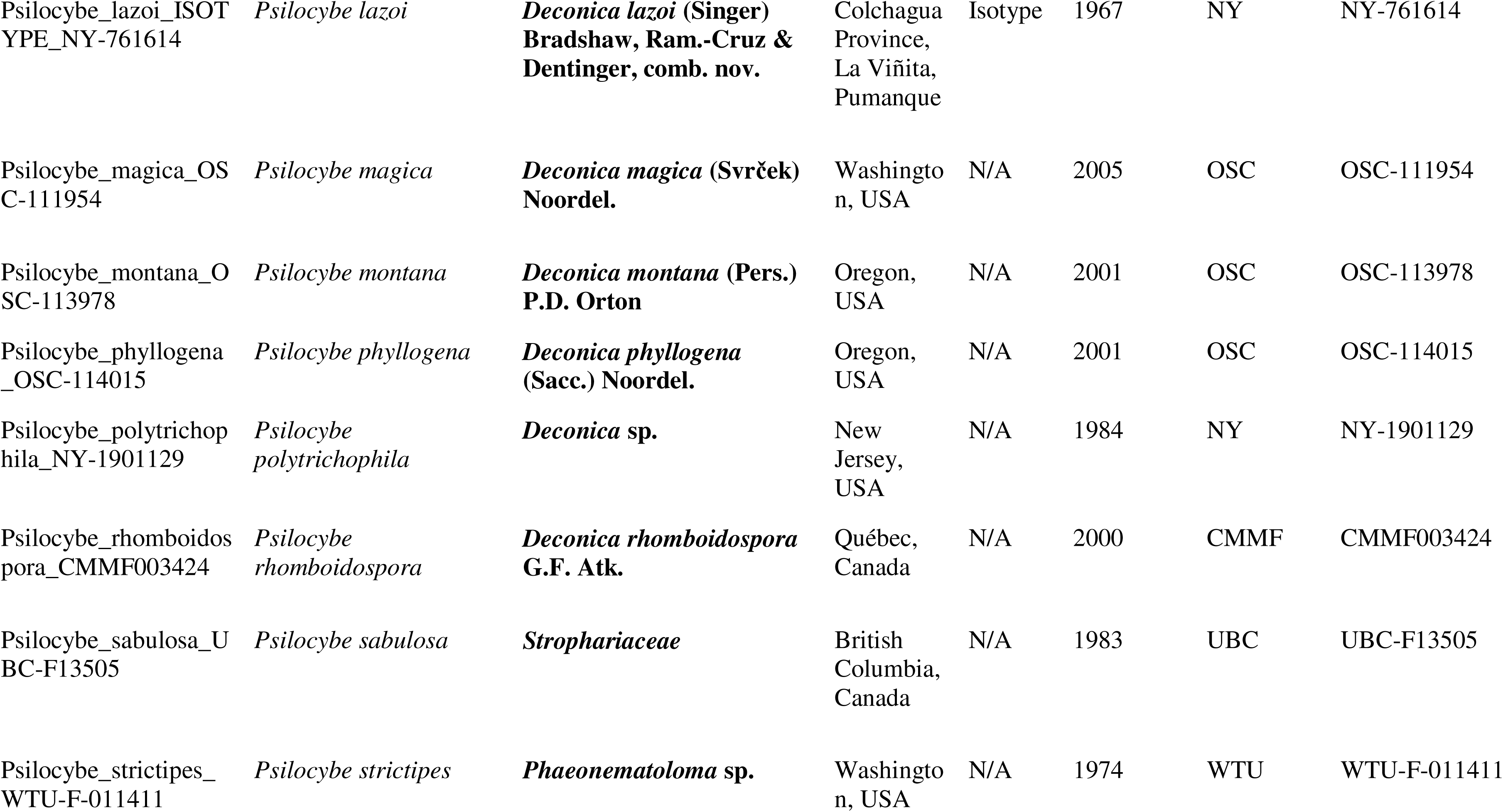
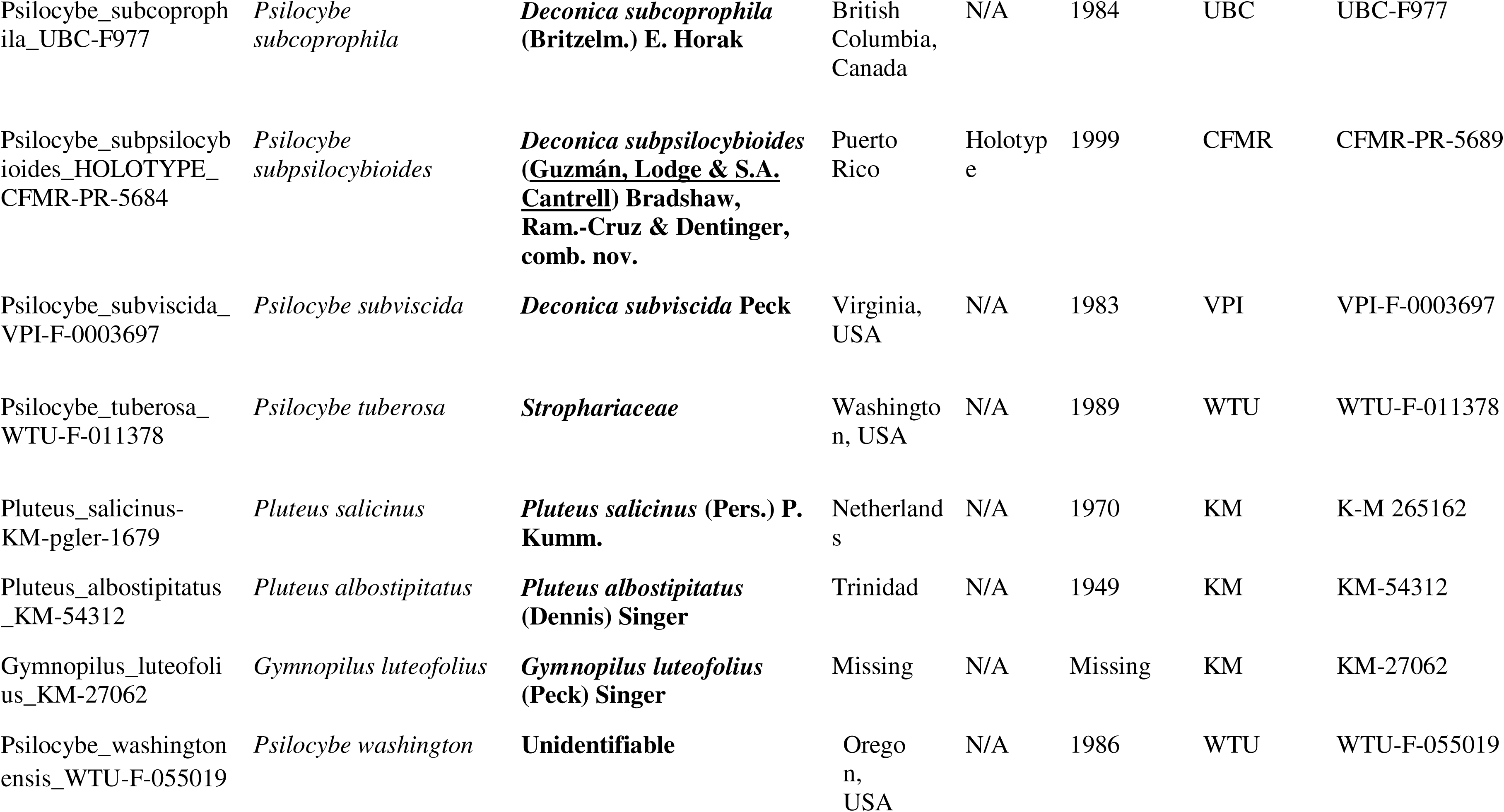
Voucher table of Specimens. Specimen voucher table for each sequenced specimen, including holding institution, catalog number, post sequencing determination, and type status of the sample if applicable.

Phylogenomic analysis using our specimens and six outgroup taxa was performed with a profile of 2983 single-copy genes derived from MCL (Markov Cluster Algorithm) clustering (Psiser 1 comparative clustering .2497) generated through MycoCosm. Two methods were used: the first was the concatenation of genes into a single supermatrix for analysis (Supplementary Figure 1). The second was a summary coalescent model of all 2983 gene trees (Supplementary Figure 2). Both trees were topologically congruent and exhibited excellent nodal support in nearly all branches. Our species tree generated two major clades that match clade I and clade II previously reported in Ramírez-Cruz et al. (2013a), but included the representation of 39 new species, which had not been investigated in that study (Figure 2, left).

**Figure 2:**
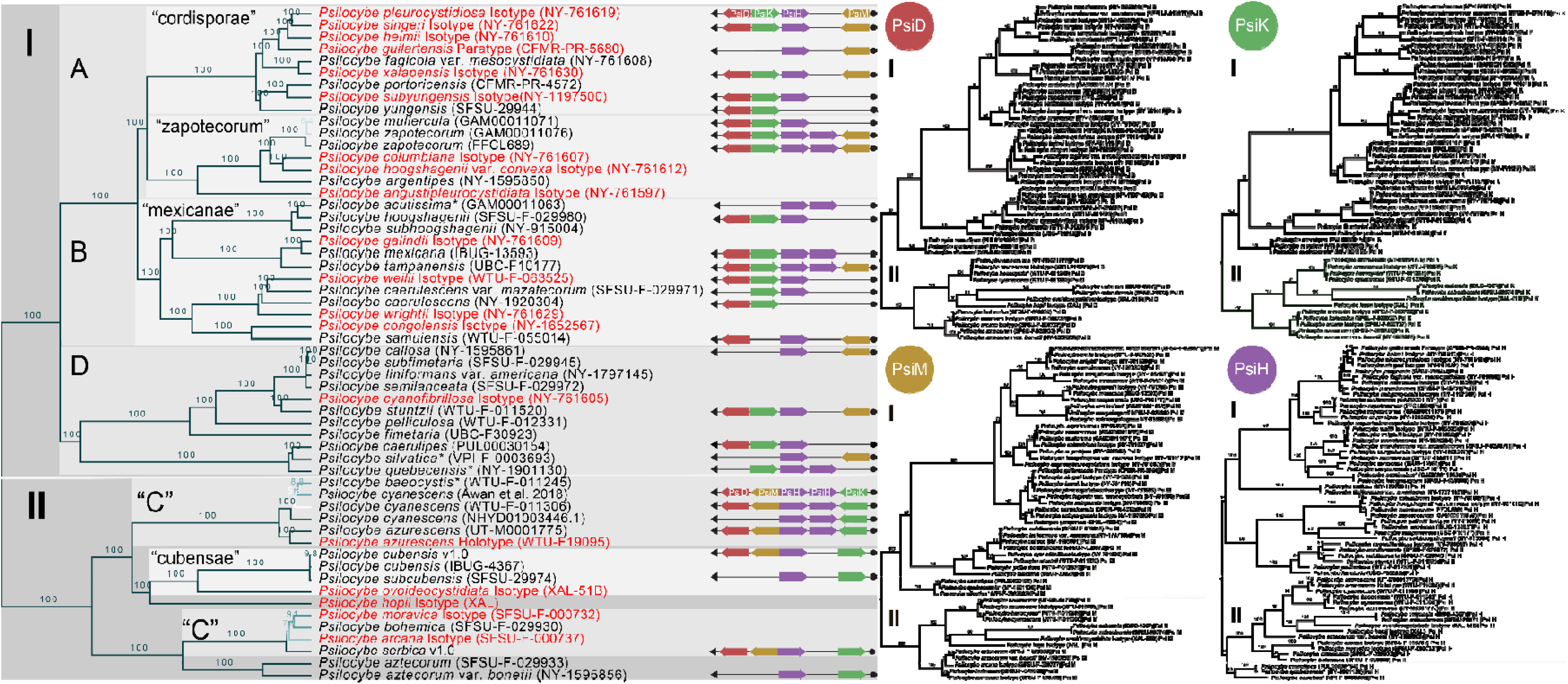
*Psilocybe* species tree, psilocybin producing gene cluster order, and Individual phylogenetic trees of the four core psilocybin producing cluster genes.

**Left:** Maximum likelihood tree constructed from 2983 concatenated genes for *Psilocybe* vouchers with major groupings of *Psilocybe* labeled and type specimens in Red. Here we expand the known major groups, while also showing that group “C” is split, which is novel to this study. **Center:** Recovered psilocybin-producing genes along with order and orientation within their respective sample. **Right:** Maximum-likelihood trees for each of the four core genes from the psilocybin producing gene cluster, PsiD, PsiK, PsiM, and PsiH. Asterisks denote vouchers determined to be misidentified.

To investigate the splitting of these major clades further, we performed molecular dating using two separate methods with calibration points set between 57 and 71 mya to represent the divergence of the family *Hymenogastraceae* (to which *Psilocybe* belongs). The first method utilized uncorrelated penalized likelihood with a “relaxed” molecular clock model, which assigned the most recent common ancestral node for clade I and clade II an age of 64.2 mya. The second method used was the estimation of divergence dates in the absence of a molecular clock through r8s (Sanderson 2003), which assigned the most recent common ancestral node for clade I and clade II an age of 60.5 mya.

### Phylogenetic analysis of ITS, EF1a, RPB1, and RPB2 genes, and macro-and micromorphology for specimen misidentification

In addition to phylogenomics, we chose to extract the universal fungal barcode, ITS (Schoch et al. 2012), and the common phylogenetic single-copy protein-coding genes elongation factor 1-alpha (EF1a), the largest subunit of RNA polymerase II (RPB1), and the second-largest subunit of RNA polymerase II (RPB2)(Supplementary Figures 3–6). We did this to rationalize our dataset with publicly available data to provide the most accurate representation of *Psilocybe* possible. Across all publicly available sequences, 40 unique taxonomic names were represented, ∼13% of the approximately 150 currently accepted *Psilocybe* s.s. However, sequence number and taxonomic representation of molecular markers varied widely. No species retrieved from the public databases had all gene regions representing a single voucher specimen. In total, the analysis of sequences from the public database along with markers extracted from our genome assemblies provided a representation of 67 unique taxa, 17 of which were represented only in the public databases. In comparison, our data set provided sequence representation for 27 taxa not previously reported. In particular, the inclusion of public data was vital for our work as it allowed us to pinpoint specimens that are likely misidentified at the species level, a common issue in museum fungaria and *Psilocybe* in particular (Figure 2, Table 1) (Andrew et al. 2019; Bradshaw et al. 2022).

While the representation of sequences for the markers was modest (EF1a n=32, RPB1 n=30, RPB2 n=15, and ITS n=26), the addition of these sequences to our phylogenomic tree enabled us to add representation for six species from EF1a, nine from RPB1, three from RPB2, and 11 from ITS for which we did not have specimens. In all gene trees, four specimen vouchers were consistently incongruent with the other taxa they clustered with: “*Psilocybe acutissima*” (GAM-00011063), “*Psilocybe baeocystis*” (WTU-F-011245), “*Psilocybe quebecensis*” (NY- 1901130), and “*Psilocybe silvatica*” (VPI-F-0003693) (Supplementary Figures 3–6). These specimens were then further analyzed using macro- and micromorphological traits to confirm that they were indeed misidentified vouchers. From the comparison of macro- and microscopic features, it was determined that GAM00011063 was mostly likely *Psilocybe hoogshagenii* R. Heim, VPI-F-003693 and NY-1901130 were mostly likely *Psilocybe caerulipes*, while WTU-F- 011245 was mostly likely *Psilocybe cyanescens* Sacc..

### New combinations in *Deconica*

Based on genomic data of type specimens, three new combinations in *Deconica* are proposed for those species that formerly were described as *Psilocybe*, in order to use the correct name.

*Deconica subpsilocybioides* (Guzmán, Lodge & S.A. Cantrell) Bradshaw, Ram.-Cruz & Dentinger, **comb. nov.**

MycoBank no.:489154

Index Fungorum Registration Identifier 489154

Basionym. – *Psilocybe subpsilocybioides* Guzmán, Lodge & S.A. Cantrell, in Guzmán, Tapia,

Ramírez-Guillén, Baroni, Lodge, Cantrell & Nieves-Rivera, Mycologia 95(6): 1174 (2003)

*Deconica clavata* (Guzmán) Bradshaw, Ram.-Cruz & Dentinger, **comb. nov.**

MycoBank no.:109209

Index Fungorum Registration Identifier 109209

Basionym. – *Psilocybe clavata* Guzmán [as ’clavatum’], Beih. Nova Hedwigia 74: 307 (1983)

*Deconica lazoi* (Singer) Bradshaw, Ram.-Cruz & Dentinger, **comb. nov.**

MycoBank no.:337850

Index Fungorum Registration Identifier 337850

Basionym. – *Psilocybe lazoi* Singer, Beih. Nova Hedwigia 29: 242 (1969)

### Evolution of the psilocybin-producing gene cluster within *Psilocybe*

Gene prediction using Augustus and a reciprocal best blast (RBB) hit method were used to extract and confirm each of the core gene sequences (PsiD, PsiK, PsiM, and PsiH). Identifications were compared to gene prediction output to determine the order and orientation of each gene in the cluster based on the strand position (+/-) of the start codon for each gene (Figure 2, center). In addition to Augustus and RBB, we also utilized Exonerates protein2genome function to confirm our results. In most cases, the gene predictions from Exonerate matched those from Augustus/RBB, with disagreements still having correct gene hits. In a few cases, we found that Exonerate identified genes more accurately and with less ambiguity compared to our RBB method (Supplementary Table 2).

We found variability in the order of the four core psilocybin-producing cluster genes within our expanded representation of *Psilocybe* species, identifying two distinct patterns. The first pattern followed a gene order of PsiD>PsiM,>PsiH>PsiK, the canonical gene order originally reported from *Psilocybe cubensis* and *Psilocybe serbica*, and was represented entirely by clade II of our dataset. The second was PsiD>PsiK>PsiH>PsiM, found in clade I of our species tree (Figure 2, right). Across our phylogeny, the original pattern was represented by 17 *Psilocybe* specimens, while 35 of our *Psilocybe* specimens exhibited the newly identified pattern.

In addition to gene order and orientation, we investigated the molecular sequence evolution of each gene separately. Phylogenetic analysis was performed without using our species constraint tree for each core psilocybin-producing genes, PsiD, PsiK, PsiM, and PsiH (Figure 2, right). We found that PsiD, PsiK, and PsiM all had the same sample clustering and topology as our species tree, suggesting a vertical inheritance pattern for these genes in *Psilocybe*. However, the gene tree for PsiH had clustering patterns consistent with our species tree but an incongruent topology, with clade II derived within clade I PsiH, rendering clade I PsiH paraphyletic. We found PsiH was far more diverse than the other core genes upon more in-depth investigation, with a much larger number of similar genes. Further, multiple samples, primarily from clade II, were found to have multiple copies of PsiH from our gene predictions (Figure 2, right, and Supplementary Table 2).

### Psilocybin homologs genes present outside of *Psilocybe* and ecological niche patterns

In contrast to only looking at the molecular evolution of the psilocybin gene cluster within *Psilocybe*, we investigated the known horizontal gene transfer (HGT) events from *Panaeolus* (Fr.) Quél., *Gymnopilus* P. Karst., and *Pluteus* Fr., using publicly available data and newly sequenced specimens vouchered as *Gymnopilus luteofolius* (Peck) Singer, *Pluteus albostipitatus* (Dennis) Singer, and *Pluteus salicinus* (Pers.) P. Kumm (Table 1 and Figure 3, left). Gene prediction and RBB analysis yielded representation of the four core psilocybin-producing genes from the published genomes of *Panaeolus cyanescens* and *Gymnopilus dilepis* (Berk. & Broome) Singer, and our newly sequenced *Gymnopilus luteofolius* and *Pluteus salicinus* genomes (Supplementary Table 2).

**Fig 3:**
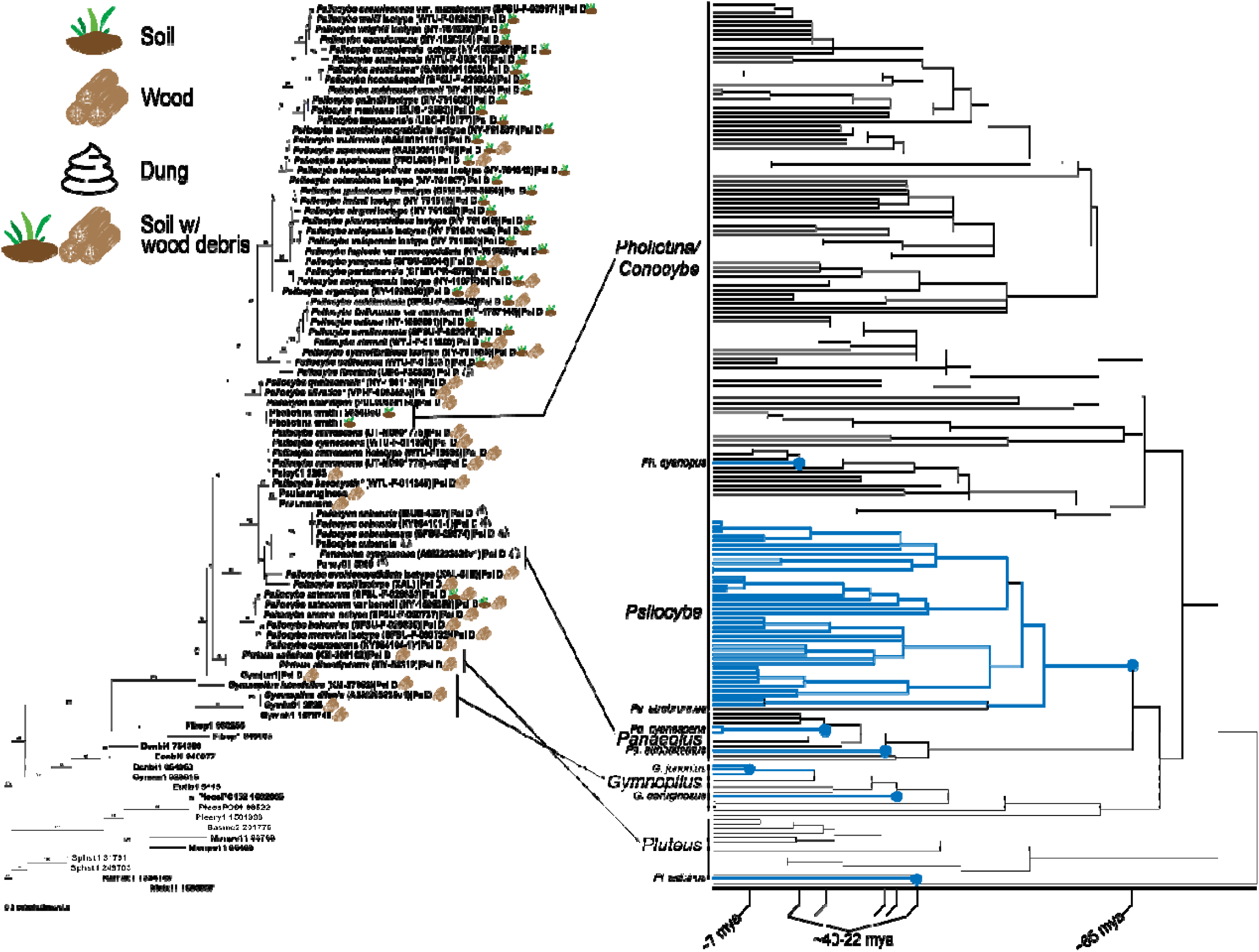
PsiD and associated ecological niches compared to psilocybin producing phylogenic analysis. **Left:** PsiD tree with associated ecological niches for *Psilocybe*, *Conocybe*/*Pholiotina*, *Pluteus*, *Gymnopilus*, and *Panaeolus,* ecological niches are represented symbolically next to sample label. **Right:** LSU phylogenetic tree of known psilocybin producing taxa, blue coloring and axis numbering correspond to psilocybin acquisitions. Asterisks denote vouchers determined to be misidentified.

Phylogenetic analysis of the PsiD gene, including those from our *Psilocybe* genomic samples, and publicly available sequences, revealed three branching patterns for our non- *Psilocybe* taxa. The first branch contained *Conocybe smithii* Watling (= *Pholiotina smithii* (Watling) Enderle) as a monophyletic branch sister to our *Psilocybe caerulipes* (Peck) Sacc. samples (BS 100%). The second branch placed *Panaeolus cyanescens* as a sister group to *Psilocybe cubensis* (BS 100%), and the final contained species of both *Gymnopilus* and *Pluteus* sharing a most recent common ancestor with all of *Psilocybe* (BS 100%) (Figure 3, left).

After expanding *Psilocybe* diversity and refining the placement of known HGT events, we sought to identify whether phylogenetic clusters could be attributed to specific ecological niches. Interestingly, our two primary phylogenetic clades correspond to distinct ecological lifestyles, with a few notable exceptions (Figure 3, left). The clade I cluster, identified through our PsiD analysis, corresponds almost entirely to the soil-dwelling saprotrophic lifestyle, except for the *Psilocybe caerulipes* grouping, which exhibits wood decay (i.e., rotting wood rather than soil enriched with woody debris) ecology. *Psilocybe* Clade II, *Gymnopilus* and *Pluteus* were associated primarily with wood decay, and in the case of *Psilocybe cubensis* and *Panaeolus cyanescens*, a coprophilous lifestyle (Figure 3, left).

### Divergence times of taxa known to produce psilocybin compared to the molecular evolution of the decarboxylase (PsiD) gene

To further investigate the evolutionary history of psilocybin acquisition, we performed a phylogenetic analysis of taxa known to contain the psilocybin gene cluster. We utilized the large ribosomal subunit (LSU) for analysis as it had the greatest diversity representation from public resources across all the taxa studied, compared to other standard protein-coding markers such as EF1a, RPB1, and RPB2. In our analysis, we targeted six specific points of psilocybin production: 1) emergence of the *Psilocybe* genus, 2) presence in *Pluteus*, 3–4) acquisitions within *Panaeolus*, 5) acquisition in *Conocybe* Fayod/*Pholiotina* Fayod, and 6) presence within *Gymnopilus* (Figure 3, right). Divergence dating of these taxa suggests that *Psilocybe* is the oldest lineage to produce psilocybin, having diverged between ∼50 mya and ∼68 mya, similar to the divergence times observed from our phylogenomic dataset of ∼65 mya. Next, presence in *Pluteus* emerged between ∼40 mya and ∼53 mya. In *Panaeolus*, phylogenetic analysis showed two, possibly individual, acquisitions for *Panaeolus cyanescens* and *P. subbalteatu*s occurring between ∼15-21 mya and ∼25-43 mya, respectively. Further, acquisition in *Conocybe*/*Pholiotina* seems to have happened between ∼26 mya and ∼36 mya, and finally the most recent emergence in *Gymnopilus* between ∼6 and 9 mya (Supplementary Table 3).

Comparison of the taxonomic relationship of *Psilocybe* and non-*Psilocybe* psilocybin producing taxa to the molecular evolution of PsiD for these same groups reveals that psilocybin likely emerged first in *Psilocybe*, which has passed the gene cluster to *Panaeolus* and *Conocybe*/*Pholiotina*. However, PsiD genes for both *Gymnopilus* and *Pluteus* are more closely related to one another than to *Psilocybe*. The emergence of psilocybin production in *Pluteus* occurs 30-40 my before the presence in *Gymnopilus* suggesting possible transfer from *Pluteus* to *Gymnopilus*, but more diverse representation in both groups would be needed to further refine this pattern.

## Discussion

### Phylogenomics of *Psilocybe*

Our phylogenomic analysis of 2,983 single-copy genes resulted in a single, unambiguous and statistically supported phylogeny for 52 species of *Psilocybe* s.s. This more than doubles the number of species included in previous analyses. Our focus on generating whole genomes from types, as old as 65 y, provided unequivocal placement of known species and a rich resource for authentication of modern and future samples. The perfect congruence of the topologies inferred from a partitioned supermatrix analysis and a summary coalescent of individual gene trees is strong evidence that the inferred phylogeny is robust to methodological approach. Because we utilized whole genome sequencing instead of reduced representation markers, this allowed us to explore genomic patterns within a robust phylogenetic context, illuminating a novel phylogenetic dichotomy in the arrangement of the genes that make up the psilocybin biosynthetic gene cluster (BGC). This dichotomy aligns with a deep divergence within the genus around 53 mya, providing new insights into the evolutionary history of this enigmatic and iconic natural product and a resource for potential development of new therapeutics.

Our phylogeny has nearly perfect (100%) nodal support at every branch (Fig. 1), including deeper branches within the backbone that are often difficult to resolve in mushrooms (Dentinger et al. 2016, 2016; Matheny et al. 2006; Moncalvo et al. 2000, 2002). The two main clades within *Psilocybe* as reported by Ramírez-Cruz et al. (2013a) were also recovered here, indicating an early split within the genus that coincides with the genic arrangement of the psilocybin BGC, but precedes a shift in ecology. Our results suggest that *Psilocybe* arose as primarily a wood-decomposing group that transitioned to soil after the split, with two shifts to dung-decomposition. However, critical species, such as the African dung-dwelling *P. natalensis* Gartz, D.A. Reid, M.T. Sm. & Eicker, are still missing, which may have an impact on the patterns seen here.

### New insights on *Psilocybe* taxonomy

The inclusion of taxa that had not been previously sampled reveals new infrageneric divisions of *Psilocybe* that contradict morphological-based classifications. For example, our results show that *P*. *aztecorum* R. Heim is sister to the *P*. *serbica* complex (sect. *Semilanceatae* Guzmán) rather than *P. baeocystis* in sect. *Aztecorum* Guzmán (1983). Together this group occupies a strictly temperate distribution in Europe, the USA, and high elevations of central Mexico. Also, the addition of the isotypes of *P. guilartensis* Guzmán, F. Tapia & Nieves-Riv., *P. heimii* Guzmán, *P. pleurocystidiosa* Guzmán, and *P. singeri* Guzmán, (sect. *Brunneocystidiatae* Guzmán), all belong to an expanded “cordisporae” clade. All of these species have rhomboid basidiospores (heart-shaped and hence the term “cordisporae”) and yellowish basal mycelia, confirming the morphological cohesion first proposed by Ramírez-Cruz et al. (2013a). The “zapotecorum” clade was also expanded to include *P. angustipleurocystidiata* Singer & A.H. Sm., *P. argentipes* K. Yokoy. (=*P. subcaerulipes* Hongo), *P. columbiana* Guzmán, and *P. hoogshagenii* var. *convexa* Guzmán. All of these taxa have small, ellipsoid or subrhomboid basidiospores, but otherwise unifying morphological characteristics are unknown.

Our results also provide new insights into well-known species complexes. For example, the species complex surrounding the European *P. moravica* Borov. (consisting of *P. arcana* Borov. & Hlavá ek, *P. bohemica* Šebek, *P. moravica*, and *P. serbica*) were extremely similar with two separate clustering patterns (*P. arcana*/*P. moravica* and *P. serbica*/*P. bohemica*)(Figure 2). This strongly implies they are all part of a single, variable species as proposed by Borovička et al. (2011), or undergoing rapid diversification. Similarly, *P. semilanceata* (Fr.) P. Kumm. clustered with *P. liniformans* var. *americana* Guzmán & Bas, sister to *P. callosa* (Fr.) Gillet and *P. subfimetaria* Guzmán & A.H. Sm. Although none of these were derived from types, this suggests that *P. liniformans* var. *americana* is synonymous with *P. semilanceata*, which is very close to *P. callosa* and *P. subfimetaria* (which may also be conspecific). Our results are also inconsistent with synonymies based only on morphological features. Cortés-Pérez et al. (2020) synonymized *P. weilii* Guzmán, Stamets & F. Tapia and *P. wrightii* Guzmán with *P. caerulescens* Murrill, whereas our results indicate there are substantial genetic differences between them and there is a strong possibility that, in fact, they represent distinct species.

Two very similar species, *Psilocybe cubensis* and *P. subcubensis* Guzmán, have variably been treated as one or two species. *Psilocybe subcubensis* was first segregated from *P. cubensis* in 1978 based on its smaller basidiospores and more tropical distribution (Guzmán and Vergeer 1978; Guzmán 1983). Both taxa form a species complex with very few diverging phenotypic differences between them (Alban-J. et al. 2021; Chethana et al. 2021). However, our data show a well-supported (BS 99%) split between groupings of *P. cubensis* and *P. subcubensis*, albeit with very short branches, suggesting that *P. subcubensis* is not conspecific with *P. cubensis*. The difference in spore size between these two taxa is possibly an early example of a morphological trait that indicates species divergence within the *P. cubensis* species complex.

### Evolution of Psilocybe cubensis

Interestingly, the branch leading to *P. cubensis/subcubensis* (sect. *Cubensae* sensu Guzmán 1983) is longer than expected for recently diverged taxa (Parks and Goldman 2014), suggesting that additional closely related taxa are missing (Ramírez-Cruz et al. 2013a) or the lineage has experienced accelerated evolution. Guzmán (1983) predicted that *P. cubensis*, a cow and horse-dung specialist originally described from Cuba by Earle in 1909, was brought to the Americas from Africa by the Spanish. Thus, it is possible that *P. cubensis* is a “self-domesticated” species that has undergone parallel evolution during the domestication of cattle. Although *P. cubensis* has not been officially reported from Africa, Guzmán predicted that other species close to it remain to be discovered there. The only African species included in a phylogenetic dataset is the *P. congolensis* Guzmán, S.C. Nixon & Cortés-Pérez we included, which is not closely related to *P. cubensis* (Figure 2). In contrast, Ramírez-Cruz et al. (2013a) reported that the sister taxon to *P. cubensis* is *P. thaiaerugineomaculans* Guzmán, Karun. & Ram.-Guill. from Thailand, which we also recovered in our RPB1 tree (Supplementary Figure 5). We additionally recovered *P. chuxiongensis* T. Ma & H.D. Hyde from Yunnan, China (Ma et al. 2014) as a sister taxon to *P. cubensis* in our ITS tree (Supplementary Figure 3. These results suggest an Asian rather than African origin for the progenitor of *P. cubensis*. However, sequences labeled as *P. natalensis* (KwaZulu-Natal, South Africa) deposited in NCBI do show close relationships with *P. cubensis* (Bradshaw et al. 2022), although these sequences do not have voucher information included, rendering them dubious. Nonetheless, these results align with African *Psilocybe* assigned morphologically to the Cubensae complex by Froese et al. (2016). Inclusion of types of the African species as well as new, targeted fieldwork is needed to clarify the relationship of *P. cubensis* and provide clearer insight into its origin.

### Evolution of the psilocybin biosynthetic gene cluster within *Psilocybe*

Our results are consistent with a strictly vertical inheritance pattern of the psilocybin BGC within *Psilocybe*. However, the two distinct gene order patterns correlate with a deep divergence within the genus approximately 53 mya. While it is unclear what the origin of these two patterns is, the early divergence and subsequent maintenance of the gene orders within the two clades may indicate additional variation early in the evolution of the psilocybin BGC. Such variation may have existed in the nascent stages of psilocybin biosynthesis, possibly in response to selective forces acting to shape its function. This discovery also indicates that transcriptional regulation of the genes may vary among taxa, resulting in taxon-specific metabolic profiles (Hoffmeister and Keller 2007; Yin and Keller 2011). Such knowledge could be harnessed in future bioengineering work to optimize and customize psilocybin biosynthesis, or as yet unknown analogs, in heterologous systems.

### Duplication of PsiH

Upon deeper investigation of PsiH, we found that it had considerably more sequence variation than any of the other core genes and has undergone multiple duplications or multiple losses (Figure 2, Supplementary Table 2). However, upon adding second best hit identified duplicated sequences similar to PsiH gene predictions, we found that the splitting of clade II became less severe, suggesting multiple paralogs across are present in many species of *Psilocybe*, despite having bioinformatically reducing our genomes to haploid values. These findings may be of particular importance for future uses in the pharmaceutical production of these compounds by providing multiple options of genes for optimizing the expression and production in organisms such as *Escherichia coli* Escherich and *Saccharomyces cerevisiae* (Desm.) Meyen (Adams et al. 2019; Milne et al. 2020).

While we primarily found duplications to occur in clade II taxa, we do not rule out that this pattern is far more pervasive across the genus. We utilized a sliding window analysis to build the most consecutive cluster placement possible. Due to this methodology, we can confidently report cluster gene order and orientation when genes were assembled within the same contig; however, this method does not perform as well on samples with more discontiguous assemblies, which is a hallmark of metagenomic assemblies from museum specimens. While this is the largest dataset of this kind in *Psilocybe,* future studies should place considerable effort on re-collecting and sequencing highly contiguous genomes of these specimens to continue to expand our repertoire of questions in *Psilocybe*.

### Patterns of inheritance and relative timing of psilocybin biosynthesis

Psilocybin and psilocin production has been reported in taxa evolutionarily distant from *Psilocybe*, including *Conocybe/Pholiotina*, *Gymnopilus*, *Inocybe* (Fr.) Fr., *Panaeolus*, and *Pluteus* (Christiansen et al. 1984; Gartz 1987; Guzmán et al. 1998; Awan et al. 2018). Reynolds et al. (2018) and Awan et al. (2018) showed that the psilocybin BGC was horizontally acquired in *Conocybe/Pholiotina*, *Gymnopilus*, *Panaeolus*, and *Pluteus*, but the direction and relative timing remained ambiguous. Using a secondary calibration from He et al. (2019) for the origin of Agaricales, our results show that psilocybin biosynthesis was first acquired by *Psilocybe* around 65 mya and was acquired by *Conocybe/Pholiotina*, *Gymnopilus*, *Panaeolus*, and *Pluteus* in multiple events between 44–7 mya (Figure 3, right).

In our analysis of the PsiD gene, we identified two clear HGT events from *Psilocybe* to other genera consistent with Reynolds et al. (2018) (Fig. 3): 1) *Conocybe smithii* (= *Pholiotina smithii*) is nested within *Psilocybe*, sharing a most recent common ancestor with the *Psilocybe caerulipes* complex, and 2) *Panaeolus cyanescens* is recovered sister to *Psilocybe cubensis*. However, *Gymnopilus* and *Pluteus* PsiD are not nested within *Psilocybe* (Figure 3, right). Such a pattern is inconsistent with HGT from *Psilocybe*, especially given the chronology of psilocybin evolution (Figure 3, right). Instead, the phylogeny indicates that the psilocybin BGC may have been acquired independently by *Gymnopilus*, *Pluteus*, and *Psilocybe*, from an as yet unidentified source or multiple sources. Reynolds et al. (2018) speculated that the psilocybin BGC may have originated in *Fibularhizoctonia* G.C. Adams & Kropp, a genus of anamorphic *Athelia* Pers. that has multiple copies of Psi homologs, albeit not contained in a cluster (Konkel et al. 2021). The Atheliales are close relatives of the Agaricales with a broad range of ecologies, including termite symbionts (Matsuura 2005), plant pathogens, mycorrhizal mutualists, and saprobes. Reynolds et al. (2018) speculated that this insect association may have provided the selective force for the evolution of psilocybin as a modulator of the symbiosis. Interestingly, *Athelia arachnoidea* (Berk.) Jülich is mycoparasitic on lichens common on tree bark in north temperate regions (Arvidsson 1976, 1978), whereas its anamorph *Fibularhizoctonia carotae* (Rader) G.C. Adams & Kropp is pathogenic on carrot roots (Adams & Kropp 1996). Our results suggest wood decomposition as the ancestral ecology of *Psilocybe*, and both *Gymnopilus* and *Pluteus* are wood decomposers. This correlation of wood as a substrate for these mushrooms and the mycoparasitic ecology of the teleomorphic *Athelia arachnoidea* on bark in shared habitats is intriguing. Mycoparasitism may be one way the psilocybin BGC could be transferred horizontally and could explain the multiple HGT events to *Gymnopilus*, *Pluteus*, and *Psilocybe* inferred from our Psi gene phylogenies and that occurred as long ago as 65 mya and as recently as 7 mya. However, *Psilocybe* spp. are not known to have mycoparasitic behaviors that would explain the clear HGT events to *Panaeolus* and *Conocybe*/*Pholiotina*. That does not mean that they do not, however, as this behavior has never been examined in *Psilocybe* and other mushrooms can exhibit dual mycoparasitic and saprotrophic ecologies (Caiafa and Smith 2022).

### Insights into the functional role of psilocybin

Despite its prominence in the scientific and public imagination, the functional role of psilocybin remains an enigma. The prevailing explanation is that psilocybin functions as an insect antifungivory defense compound (Reynolds et al. 2018). However, this prevailing idea requires more investigation as psilocybin mushrooms are often reported to be occupied by insects laying eggs or living larvae in senesced field collections. Psilocybin is a prodrug that is rapidly converted to the dephorphorylated psilocin, which mimics serotonin and binds tightly to serotonin receptors, especially 5-HT_2A_, a pharmacological action common to many psychoactive tryptamines (Araújo et al. 2015). High-affinity binding to these receptors in mammals, and homologs in distantly related organisms such as insects and arachnids, produces unnatural and altered behaviors (Christiansen et al. 1962; Nichols et al. 2002; Passie et al. 2002; Boyce et al. 2019). This disorientation may be a direct deterrent or could render the subject more vulnerable to predation (or, alternatively, may be a reward). Serotonin receptors are also highly expressed in the lining of a wide range of animal gastrointestinal tracts where serotonin is involved in digestion (French et al. 2014). Thus, it is also possible that psilocybin functions as an emetic or laxative to promote the dispersal of spores before they are rendered inviable by digestion. However, other than humans, there are few documented cases of vertebrates consuming psilocybin-containing mushrooms (consisting entirely of domesticated dogs) (Beug et al. 2006), many of which are uncommon or scarce and would be difficult to learn to recognize with minimal encounters, limiting their selective potential. Moreover, anecdotes of larvae-infested *Psilocybe* abound and one experiment showed that adult flies could be reared from *P. cyanescens* mushrooms that were identical to flies reared from a co-occurring non-psilocybin containing mushroom (Awan et al. 2018). Experimental evidence for the functional role of psilocybin is thus far lacking.

Lenz et al. (2020) elucidated the chemical basis of the blue pigment that forms when psilocybin-containing mushrooms are damaged as oligo- and poly-mers of psilocin. The conversion of psilocybin to psilocin and the linking of them into chains is enzymatically controlled. Lenz et al. (2020) also pointed out that the psilocin oligo/polymers have chemical properties similar to plant flavonoids and polyphenolic tannins, which produce reactive oxygen molecules that can cause lesioning in the basic and oxidative environment of invertebrate guts. Thus, the psilocin oligo/polymers may be an inducible defense against fungivory, whereas psilocybin may function simply as the building block kept in reserve for the true chemical weaponry. Intriguingly, the formation of the blue psilocin oligo/polymers is invariably connected to psilocybin biosynthesis throughout the multiple independent inheritances (Reynolds et al. 2018; Awan et al. 2018) and convergent evolution (Awan et al. 2018). The maintenance of the enzymatic capacity to inducibly convert psilocybin lends further support to the hypothesis that it is the blue oligo/polymer with an ecological function rather than the possibly accidental pharmacological effects of psilocybin, itself.

Although empirical evidence of inducible defenses against fungivory in mushrooms is largely lacking, one study demonstrated experimentally that the ubiquitous mushroom volatile 1- octen-3-ol is an inducible slug antifeedant (Wood et al. 2000). Interestingly, 1-octen-3-ol is also an insect attractant, including mosquitoes that are attracted to it in human breath and sweat (Takken and Kline 1989). Other fungivorous insects are also known to be attracted to 1-octen-3- ol including Culicoides (midges) and tsetse flies (Hall et al. 1984; Kline et al. 1994; Grant and Dickens 2011), suggesting that its recognition is likely fairly ancient as it would indicate food sources or brooding sites. In fact, 1-octen-3-ol is a critical element of the mushroom mimicry displayed by *Dracula* Luer orchids to attract pollinating *Zygothrica* Wiedemann flies (Joulain 1993; Kaiser 2006; Policha et al. 2016). The attractive quality of 1-octen-3-ol to insects and its ubiquitous presence in mushrooms are inconsistent with the hypothesis that insect fungivory has widespread negative impacts in mushrooms. However, the slug antifeedant property of 1-octen- 3-ol suggests it may primarily function to stave off damage from fungivorous gastropods. Intriguingly, terrestrial gastropods first arose and diversified at the end of the Cretaceous mass extinction (Sohl 1987; Morris and Taylor 2000; Kaim 2002; Neubauer and Harzhauser 2022), concomitant with our inferred origin of psilocybin biosynthesis in *Psilocybe*. In contrast, insects and angiosperms underwent evolutionary expansion during the Cretaceous from 145.5 and 65.5 mya, but suffered mass extinction following the impact of the asteroid that, by some estimates, caused nearly 80% of living species to go extinct (Raup and Boyajian 1988; Briggs 1991). Taken together, we hypothesize that psilocybin evolved as a pro-building block for an inducible chemical defense against fungivorous gastropods. Experimental studies will be critical to test this hypothesis.

### Conclusion

Since the “re-discovery” of *Psilocybe*, this group of organisms and its unique psychoactivity has been of great interest from cultural, counter-cultural, scientific, and medical perspectives. However, *Psilocybe*, and the production of psilocybin and psilocin in general, hold numerous biological secrets that remain to be elucidated. Our study highlights how whole genome sequencing can enrich systematics studies by providing opportunities to examine patterns of gene evolution of a phenotypic trait. By mining genomic data beyond markers for phylogenetic inference, we gained novel insights into the evolutionary origins of psilocybin biosynthesis that have implications for understanding the functional role of this powerful chemical and can inform translational applications for human well-being.

## Methods

### DNA extraction and genomic sequencing

Sample tissue was extracted from the hymenophore ranging from 5–15 mg (our best extraction results usually came from large inputs between 10 and 15 mg). These fragments were homogenized by placing them in 2.0 mL screw-cap tubes containing a single 3.0 mm and 8 1.5 mm stainless steel beads and shaking them in a BeadBugmicrotube homogenizer (#Z763713, Sigma) for 120 seconds at a speed setting 3500 rpm. DNA extraction was performed with NEB genomic monarch kit (monarch kit T3010S) with in-house protocol alterations to use the provided tissue lysis buffer at double volume and increase the amount of wash buffer to 550 µl during the washing steps.

Samples that had been collected before 1950 (such as many of the type specimens) were processed with a phenol-chloroform DNA extraction protocol. In short, mechanical disruption of tissue was followed as previously stated, then total lysate was placed in phase lock gel tubes (Quantabio #2302820) along with 1:1 volume of phenol/chloroform solution (Calibiochem OmniPur, EMD Millipore Corporation #5.39680.0002) and then mixed for 10 minutes on a hula mixer. Homogeneous mixtures were centrifuged at maximum speed for 10 minutes, then the aqueous (top) layer was transferred to a new phase-lock gel tube, and each step was repeated a second time before moving on. After extractions, precipitation was performed by ethanol precipitation with 7.5 molar ammonium acetate. gDNA samples were submitted to the High Throughput Genomics Core at the University of Utah, where libraries were prepared using Nextera DNA Flex Library Prep (cat: 20018704) and sequenced on a full lane of Novaseq 2× 150 bp.

### Quality control and genome assembly

Sequencing run statistics and quality were visualized for each sample using Fastqc version 0.11.9 (CITATION?) and then compared to each other using MultiQC version 1.10 (Ewels et al. 2016). Library estimates were generated using the EstimateLibraryComplexity function of Picard toolkit (http://broadinstitute.github.io/picard/), while genome assembly stats were quantified using a custom Perl script. Raw sequencing reads were trimmed and quality filtered using fastP version 0.20.1 (Chen et al. 2018) and then assembled using SPAdes version 3.15.2 (Bankevich et al. 2012) with kmer values alternating every other digit between 1 and 127, inclusive.

### Homologous gene identification and phylogenetic species tree construction

Homologous genes used for phylogenetic analysis were identified by accessing publicly available MCL gene sequences from the Psiser1 comparative clustering.2497 profile (Fricke et al. 2017) through the Joint Genome Institute’s (JGI) Mycocosm genome portal (Grigoriev et al. 2012, 2014). This profile included two species of *Psilocybe* (*P. cubensis* and *P. serbica*) (Fricke et al. 2017) and six closely related taxonomic outgroups, *Agrocybe pediades* (Fr.) Fayod (Agrped1) (Ruiz-Dueñas et al. 2021), *Galerina marginata* (Batsch) Kühner (Galma1) (Riley et al. 2014), *Gymnopilus chrysopellus* (Berk. & M.A. Curtis) Murrill (Gymch1), *G. junoniu*s (Fr.) P.D. Orton (Gymjun1) (Ruiz-Dueñas et al. 2021), *Hebeloma cylindrosporum* Romagn. (Hebcy2) (Mycorrhizal Genomics Initiative Consortium et al. 2015), and *Pholiota alnicola* (Fr.) Singer (=*Flammula alnicola* (Fr.) P. Kumm.)) (Phoaln1)(Ruiz-Dueñas et al. 2021), which contained 2,983 single-copy genes. While not the closest relative of *Psilocybe*, reference sequences for these 2,983 were taken from Galma1:v1 (*Galerina marginata*) (Riley et al. 2014) to facilitate a more conservative approach to homology identification in our sequenced genomes.

The Galma1 reference sequences were then used as a basis for the pairwise alignment tool Exonerate version 2.4.0 (Guy St.C. 2000) to extract representative sequences of each single-copy gene for each sample genome. All genes were then aligned using the multiple sequence alignment program MAFFT version 7.475? (Katoh 2002) with the parameters --maxiterate 1000 --localpair --reorder. MAFFT alignments were concatenated with a custom script, which also provided a partition file for subsequent tree reconstruction. Concatenated sequences and partitions file were then used for phylogenetic through IQ-TREE version (Nguyen et al. 2015) using automatic model finder (Kalyaanamoorthy et al. 2017) and 1000 ultrafast bootstrap replicates optimized using the --bnni flag (Hoang et al. 2018).

Multi-gene summary coalescent tree analysis was conducted in the same manner as previously stated, except that each gene profile was aligned individually rather than concatenated. Each gene alignment was then used to construct a phylogenetic tree in the same manner listed previously. Coalescent analysis was performed using all 2,983 gene trees to construct a consensus tree using ASTRAL-III (Zhang et al. 2018). Both trees were congruent with one another, and the concatenated gene tree was chosen to be used as a constraint tree for further DNA barcode analysis.

### Molecular dating and divergence of phylogenomic data

Molecular divergence dating analysis was conducted on our *Deconica*, *Psilocybe*, and outgroup taxa concatenated tree (Supplementary Figure 1) utilizing two different methods with a calibration range of 71 mya (Ruiz-Dueñas et al. 2021) to 57 mya (Varga et al. 2019), representative of the divergence of the family *Hymenogastraceae*. The first method utilized uncorrelated penalized likelihood with the “relaxed” model in the R package APE version 6.6–2 (Paradis and Schliep 2019). The second method utilized was through the GUI for pyr8s version (0_1_win) (Sanderson 2003) utilizing the min-max age range calibration of 57–71 mya, with all other parameters set to default. Methods using Bayesian analysis such as BEAST2 (Bouckaert et al. 2014) were not used as the computational time needed to active convergence on a dataset of this size was unachievable. Trees and divergence times were visualized using FigTree version 1.4.4 (http://tree.bio.ed.ac.uk/software/figtree/ ).

### ITS, EF1a, RPB1, and RPB2 gene extraction and phylogenetic analysis

DNA barcodes were extracted bioinformatically with an in-house pipeline. In brief, 10859 complete fungal ITS, 487 EF1a, 639 RPB1, and 937 RPB2 with conserved exon sequences from AFTOL (Lutzoni et al. 2004) were aligned respectively with MAFFTversion 7.475 (Katoh 2002) and used to generate an HMM profile with hmmr version 3.1b2 (hmmer.org). The HMM profile for fungal ITS was used with PathRacer (SPAdes-3.15.0-pathracer-2020-12-20-dev) (Shlemov and Korobeynikov 2019) in conjunction with the assembly graph output from SPAdes to produce a most probable sequence path for the HMM provided. ITS sequences were manually trimmed from edge sequence outputs. Sequences of EF1a, RPB1, and RPB2 were extracted from edge sequence outputs with Exonerate using the flag-protein2genome and a query of GenBank proteins sequences from *Psilocybe cyanescens* (ADI71893.1- EF1a, AHB18799.1-RPB1, and AHH34099.1-RPB2). All *Psilocybe* s.s. sequences corresponding to the same loci (ITS, EF1a, RPB1, RPB2) were retrieved from NCBI as well as all species hypotheses (SH) ITS sequences from the curated fungal database UNITE (Abarenkov et al. 2010). Each data set from NCBI was checked for overlap of different genes for a single vouchered specimen. Due to the robust nodal support of our phylogenomic species tree, analysis was performed using the concatenated gene tree constructed with only *Psilocybe* specimens in conjunction with all public datasets.

Phylogenetic trees for each barcode were constructed using IQ-TREE with the same parameters as above, except for using a constraint tree derived from the concatenated gene tree to preserve the well-supported topology from the phylogenomic analysis.

### Microscopic characteristics to confirm misidentification

For microscopic investigation, hand sections were made from the pileus to observe basidiospores and cheilocystidia. Sections were mounted in 5% KOH after being rehydrated with 70% ethanol. At minimum of 25 measurements of each microstructure were taken using a 100 × oil-inmersion objective on Zeiss K-7 light microscopes and photographed using a Zeiss Axioscop 40 with Zen 2.3 lite.

### Gene prediction of psilocybin gene cluster and construction of gene trees

With robust phylogenomic data, questions outside the realm of phylogenetic species relationships can now be explored, such as the evolution of specific traits and their ecological role. In particular, the psychoactive secondary metabolites, psilocybin and psilocin, first identified in *Psilocybe* mushrooms, are of particular interest due to their high therapeutic potential (Frood 2008; Lowe et al. 2021; Marks and Cohen 2021). We used gene prediction and an RBB bioinformatic extraction method to identify each gene from the clusters in our genomes and phylogenetic analysis of each of our genes was generally congruent with our species tree (Figure 2), except for the P_450_-encoding PsiH.

Due to the diploid nature of our genomes, we first used the software package Redundans version 0.14a (Pryszcz and Gabaldón 2016) to phase our assemblies before gene prediction. Psilocybin gene cluster detection was performed using the Augustus version 3.4.0 gene prediction software (Stanke et al. 2006) with the flags –singlestrand=true to predict genes independently on each strand, and using the coprinus_cinereus training set. Following gene prediction, the most likely psilocybin BGC for each species was chosen as follows: using the protein sequences for PsiD, PsiK, PsiH, and PsiM based on the publicly available chromosomal level assembly for *P. cubensis* (McKernan et al. 2021), we used BLASTP with default parameters to search for homologs in the predicted translated proteins for each species. The top three hits for each gene were then used together to find putative clusters by examining the gene numbers assigned by Augustus. Putative “clusters” were identified as any subset of the 21 genes where the predicted gene numbers of any gene within the subset was at most two genes away from one other gene in the subset. This set of computationally-generated putative psilocybin clusters was then manually curated by querying the top three hits against the Psi gene sequences from the chromosomal assembly of *P. cubensis* using BLASTn to measure their similarities. The largest subset with high similarities to the Psi genes was chosen as the cluster candidate for each *Psilocybe* species (Figure 2). To corroborate our gene predictions, we utilized Exonerate with the same Psi protein queries. Phylogenetic trees were generated as above, without utilizing the species constraint tree, then visualized and midpoint-rooted using FigTree 1.4.4.

### PsiD molecular analysis with *Psilocybe* and non-*Psilocybe* psilocybin producers

PsiD nucleotide sequences extracted through exonerate were transcribed utilizing codon-aware multiple sequence alignment through HYPHY version 2.5.36 (Pond et al. 2005) following the tutorial outlined at: https://github.com/veg/hyphy-analyses/blob/master/codon-msa/README.md

Extracted PsiD amino acid sequences were combined with the PsiD ammino acid sequences published in Reynolds et al. (2018), undergoing multiple sequence alignment and phylogenetic analysis under the same parameters used for individual gene trees.

### Psilocybin producing taxa phylogenetic analysis and divergence dating

In order to assess the relative timing of psilocybin biosynthesis across the mushrooms, a dataset was constructed of all publicly available sequences of the ribosomal large subunit DNA region (28S; downloaded from NCBI on 14 August 2022) for *Conocybe/Pholiotina,Gymnopilus, Panaeolus, Pluteus, and Psilocybe.*. A sequence of *Calocybe gambosa* (Fr.) Donk was used as an outgroup. Due to the poor overlap between taxa for different gene regions, the 28S region had the best species representation that could be aligned across all genera. Multiple sequence alignment was performed using the L-INS-i algorithm in MAFFT. Phylogenetic analysis was performed under maximum likelihood using IQ-Tree with automatic model selection and up to 1000 nonparametric rapid bootstraps. Branch lengths were converted to relative time using uncorrelated penalized likelihood implemented in the ‘chronos’ function in the R package ‘ape’. Dates were inferred from the distance of the nodes from the tips based on the distance to the stem node for *Psilocybe* estimated, above. Additionally, pyr8s was run with a time constraint of 136 mya, corresponding to the divergence date of *Agaricales* (He et al. 2019), as well as in scalar mode for comparison (Supplementary Table 3).

## Acknowledgments

We acknowledge the Natural History Museum of Utah for its commitment to collaborative Science and the Genomics Core Facility, a part of the Health Sciences Cores at The University of Utah, for their input and high-quality work. Additionally, we acknowledge the numerous institutions that provided specimens for destructive sampling, many of which were rare and irreplaceable. Further, we wish to recognize the dedicated and hard work performed by Isabelle Galland and Talia A. Backman in their help processing many of these samples for DNA sequencing. Further, the authors wish to thank David Scott Flocken for his insight and thought- provoking conversations on the work generated here. Additionally, we would like to thank Jan Borovička and Oscar Castro-Jauregui for providing high-quality images and graciously allowing us to use them for publication. Additionally, we would like to thank Dr. Jason Slot for providing data from previous publications as well as intriguing conversions about Psilocybin evolution. VRC and LGD would like to thank the University of Guadalajara and CONACYT for their support of their research.

## Data Availability

Raw short-read sequences for this project have been deposited in the Sequence Read Archive (SRA) and assigned the bioproject number PRJNA904752 . Raw tree files, assemblies, gene prediction files, multiple sequence alignments, and *Psilocybe* constraint tree are available through Dryad under https://doi.org/10.5061/dryad.tmpg4f52s. Any code or specific script requests should be sent to the corresponding author.

## Author contributions

Alexander J Bradshaw performed all molecular processing of specimens, genome assembly, phylogenomic analysis, data analysis, and initial gene prediction in addition to preparing and writing this manuscript. Virginia Ramírez-Cruz contributed with outreach to collection institutions, microscopic confirmation of misidentified specimens, taxonomic confirmation and updating information as well as providing crucial insight into the preparation and editing of this manuscript. Ali R. Awan performed bioinformatic analysis to identify and extract the psilocybin gene cluster genes from each specimen for use in phylogenetic analysis and the preparation and writing of this manuscript. Giuliana Furci contributed with outreach to collection institutions and comments and additions to the manuscript. Laura Guzmán-Dávalos contributed with outreach to collection institutions and comments, additions, and editing of the manuscript. Paul Stamets contributed comments and additions to the manuscript, as well as high-resolution images for specimens. Bryn T.M. Dentinger provided funding, experimental, and data analysis guidance and assisted in preparing and editing this manuscript.

## Supplementary Figures

**SF 1- Concatenated phylogenetic tree with *Deconica* genomes**

**SF 2- ASTRAL-III phylogenetic species tree with *Deconica* genomes**

**SF 3- ITS phylogenetic tree using *Psilocybe* species constraint tree**

**SF 4- EF1a phylogenetic tree using *Psilocybe* species constraint tree**

**SF 5- RPB1 phylogenetic tree using *Psilocybe* species constraint tree**

**SF 6- RPB2 tree using *Psilocybe* species constraint tree**

## Supplementary Tables

**ST-1-Genome assembly and BUSCO stats for each specimen**

**ST-2- RBB gene numbers, contigs, clustering results, and RBB method comparison to Exonerate**

**ST-3- Divergence time analysis methods and results**

