## Supplementary figures and images for "Phylogenomics of the psychoactive mushroom genus *Psilocybe* and evolution of the psilocybin biosynthetic gene cluster"

### SF 1-Concatenated Phylogenetic tree with all genomes.pdf

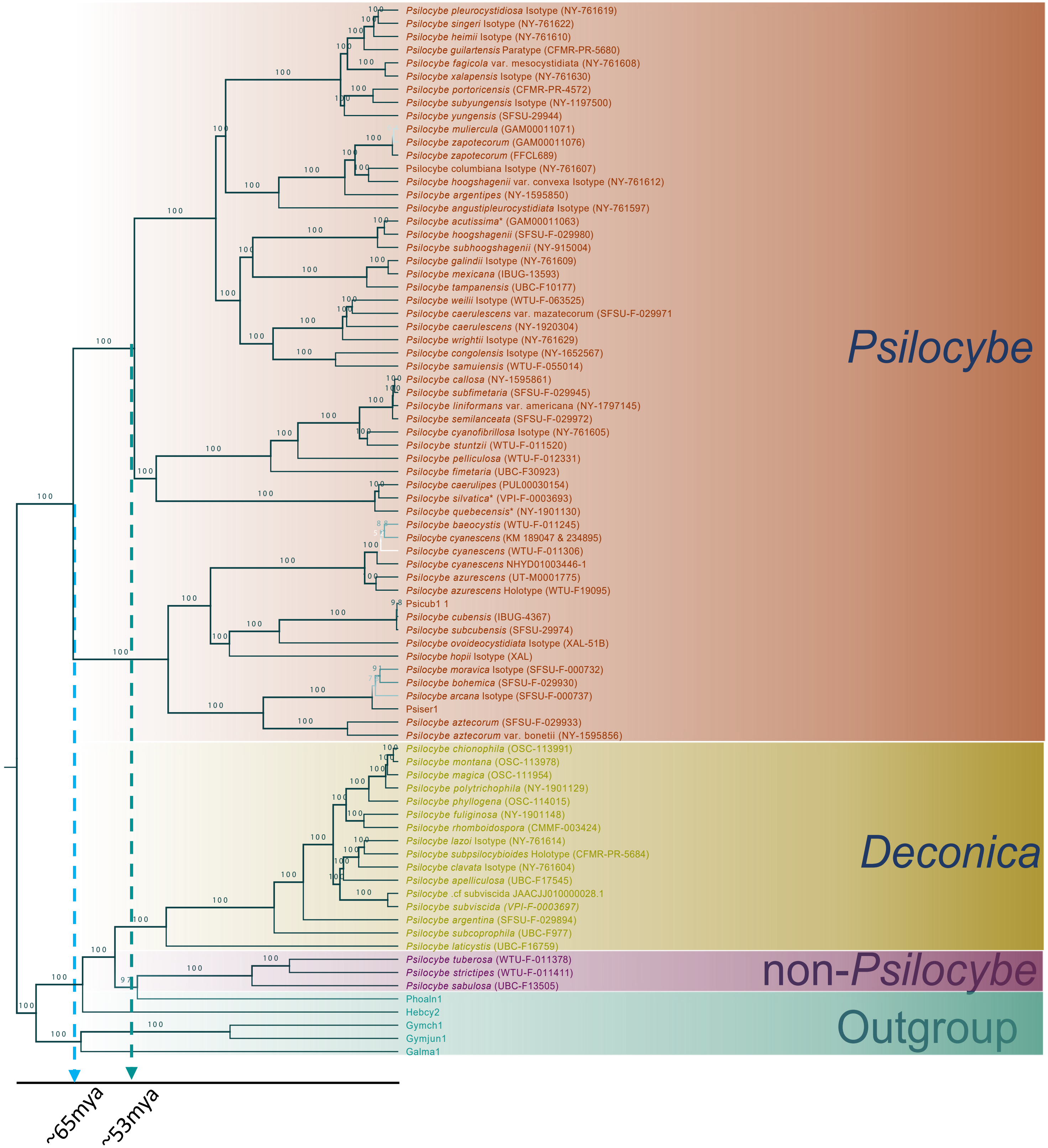

### SF 2- ASTRAL-III Phylogenetic species with Deconica genomes.pdf

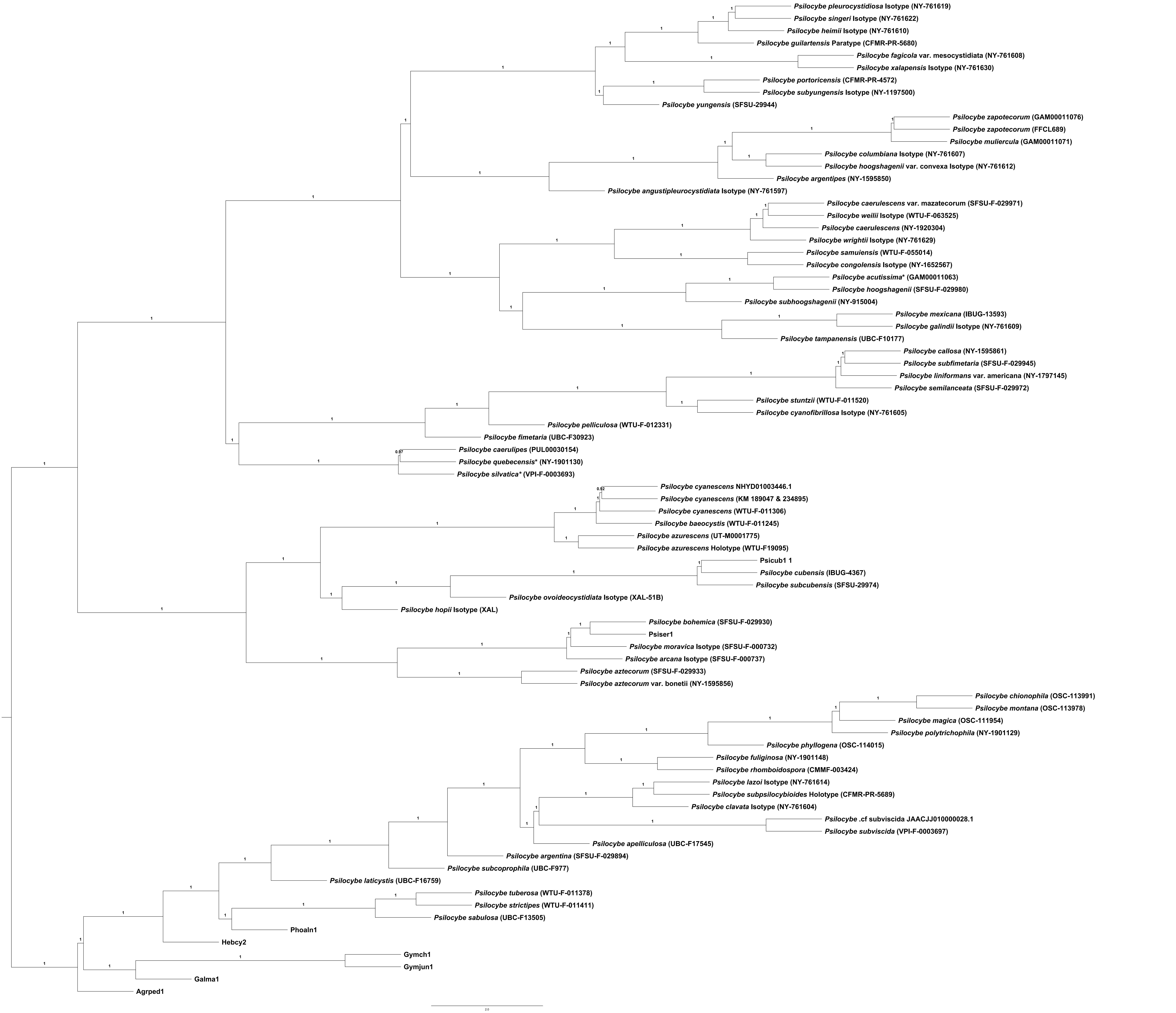

### SF 3-ITS phylogenetic tree using psilocybe constraint tree.pdf

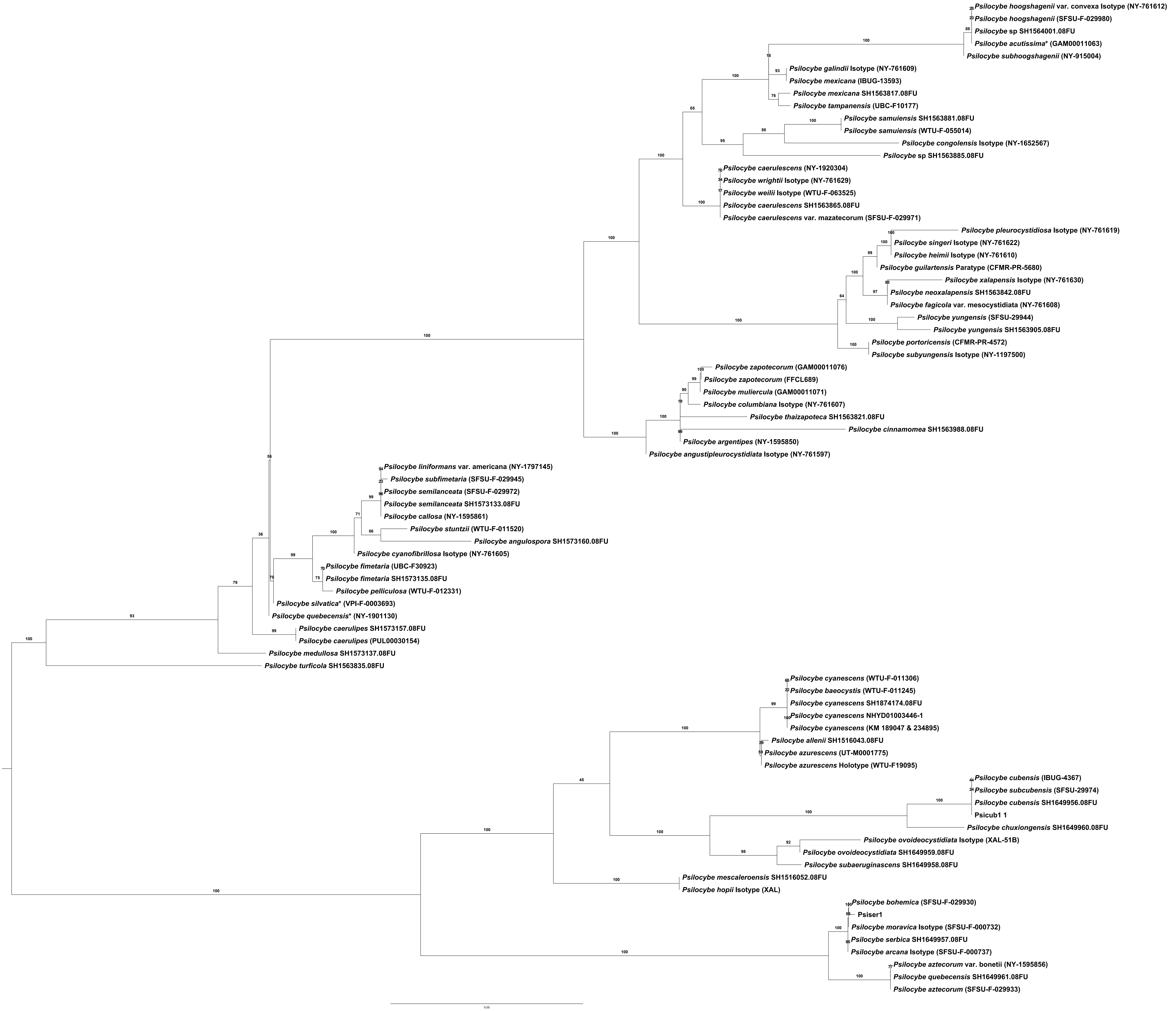

### SF 4-EF1a phylogenetic tree using phylogenomic constraint tree.pdf

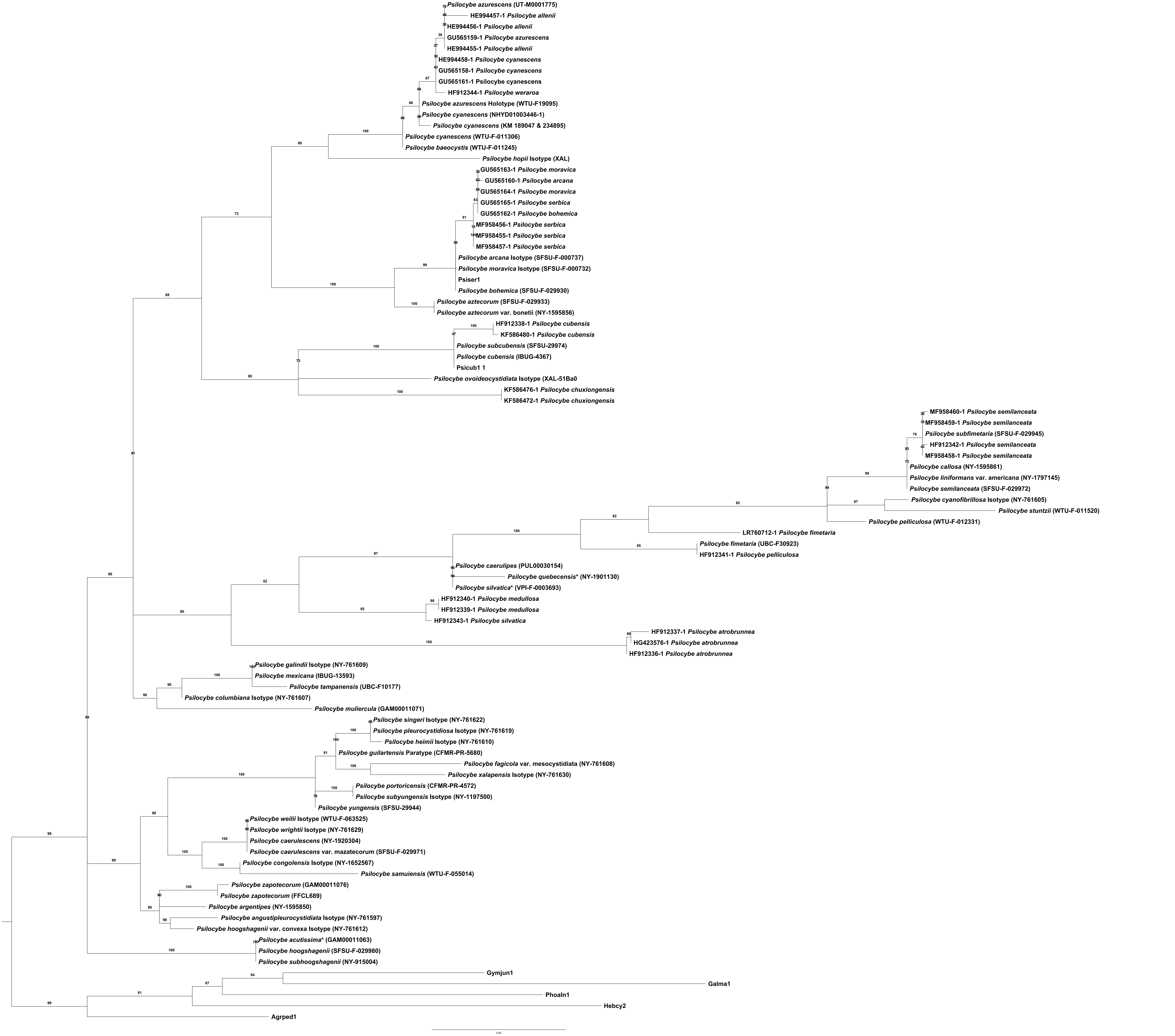

### SF 5-RPB1 phylogenetic tree using phylogenomic constraint tree.pdf

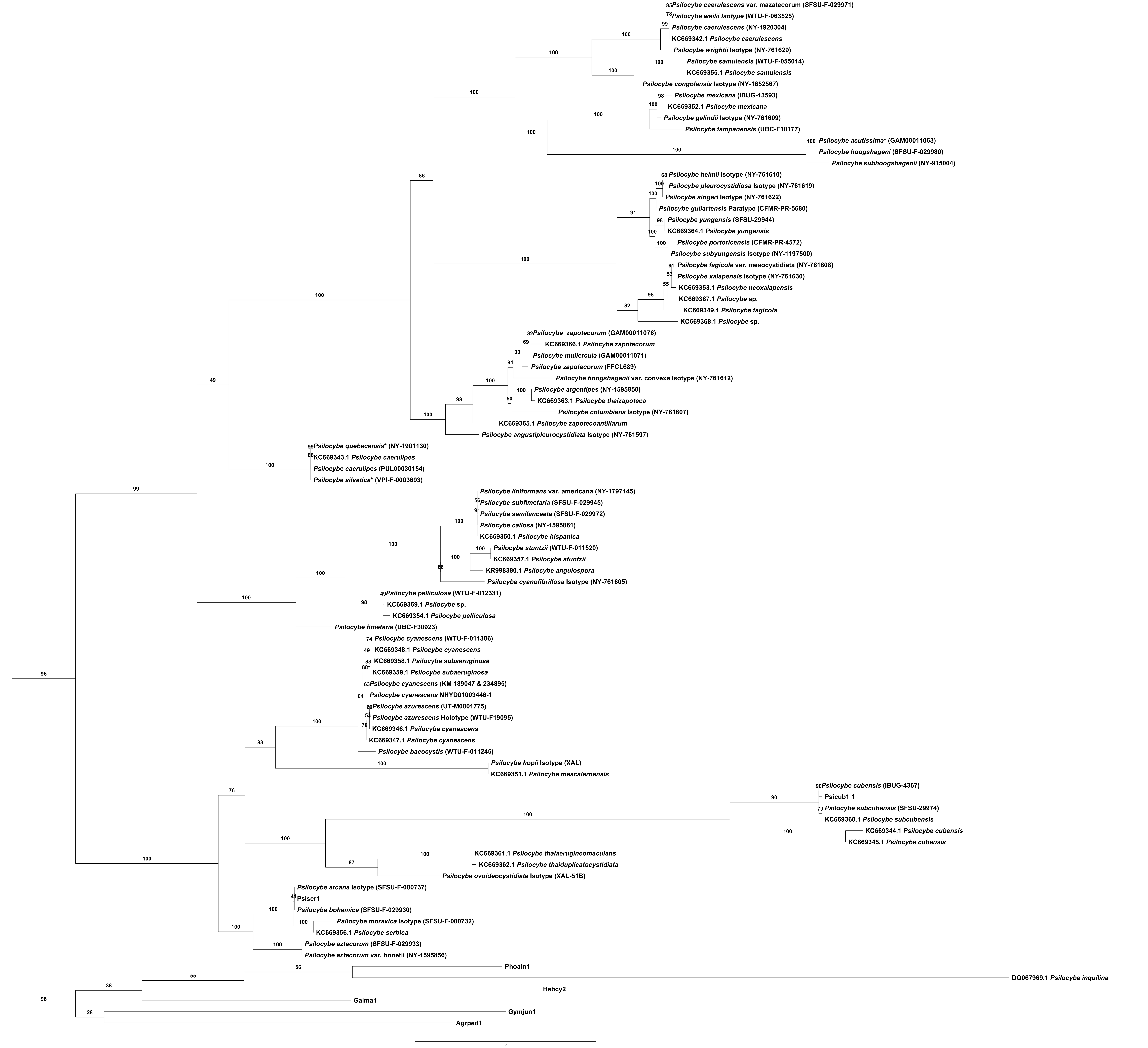

### SF 6-RPB2 phylogenetic tree using psilocybe constraint tree.pdf

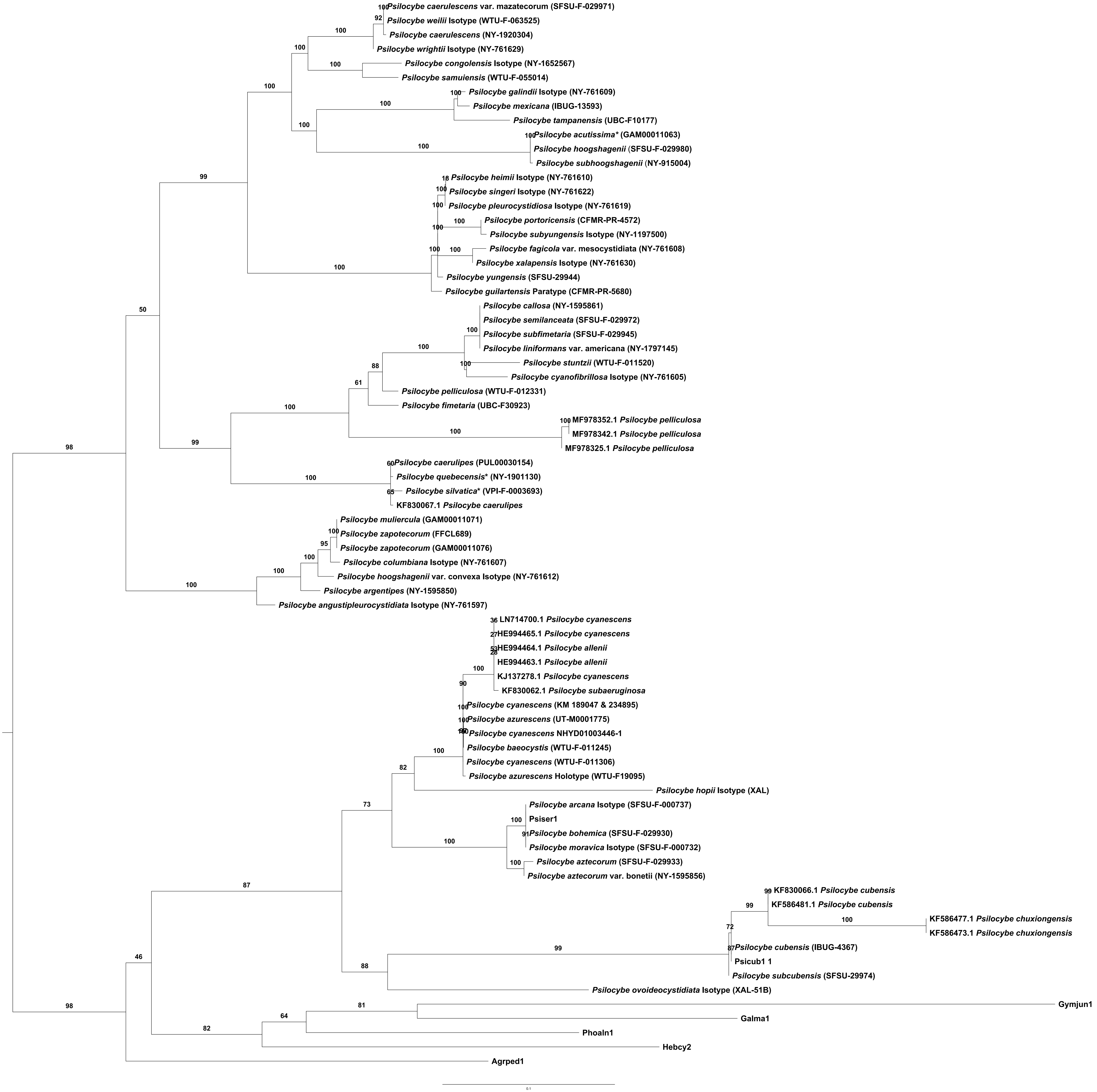
